# Metataxonomic insights in the distribution of *Lactobacillaceae* in foods and food environments

**DOI:** 10.1101/2022.09.09.507241

**Authors:** Eugenio Parente, Teresa Zotta, Marilisa Giavalisco, Annamaria Ricciardi

## Abstract

Members of the family *Lactobacillaceae*, which now includes species formerly belonging to the genera *Lactobacillus* and *Pediococcus*, but also *Leuconostocaceae*, are of foremost importance in food fermentations and spoilage, but also as components of animal and human microbiota and as potentially pathogenic microorganisms. Knowledge of the ecological distribution of a given species and genus is important, among other things, for the inclusion in lists of microorganisms with a Qualified Presumption of Safety or with beneficial use. The objective of this work is to use the data in FoodMicrobionet database to obtain quantitative insights (in terms of both abundance and prevalence) on the distribution of these bacteria in foods and food environments.

We first explored the reliability of taxonomic assignments using the SILVA v138.1 reference database with full length and partial sequences of the 16S rRNA gene for type strain sequences. Full length 16S rRNA gene sequences allow a reasonably good classification at the genus and species level in phylogenetic trees but shorter sequences (V1-V3, V3-V4, V4) perform much worse, with type strains of many species sharing identical V4 and V3-V4 sequences. Taxonomic assignment at the genus level of 16S rRNA genes sequences and the SILVA v138.1 reference database can be done for almost all genera of the family *Lactobacillaceae* with a high degree of confidence for full length sequences, and with a satisfactory level of accuracy for the V1-V3 regions. Results for the V3-V4 and V4 region are still acceptable but significantly worse. Taxonomic assignment at the species level for sequences for the V1-V3, V3-V4, V4 regions of the 16S rRNA gene of members of the family *Lactobacillaceae* is hardly possible and, even for full length sequences, and only 49.9% of the type strain sequences can be unambiguously assigned to species.

We then used the FoodMicrobionet database to evaluate the prevalence and abundance of *Lactobacillaceae* in food samples and in food related environments. Generalist and specialist genera were clearly evident. The ecological distribution of several genera was confirmed and insights on the distribution and potential origin of rare genera (*Dellaglioa, Holzapfelia, Schleiferilactobacillus*) were obtained.

We also found that combining Amplicon Sequence Variants from different studies is indeed possible, but provides little additional information, even when strict criteria are used for the filtering of sequences.

## 1. Introduction

Bacteria belonging to family *Lactobacillaceae* (which now includes *Leuconostocaceae*, Zheng et al., 2020) are of foremost importance as beneficial microorganisms in food fermentations (Anagnostopoulos and Tsaltas, 2022; Arora et al., 2020; Bourdichon et al., 2012; 2019; Gänzle, 2022; Laranjo et al., 2019; Lee et al., 2020; Mayo et al., 2021; Sumby et al., 2014; Tamang et al., 2020; Walsh et al., 2020), as components of human and animal microbiota (Duar et al., 2017; Pasolli et al., 2020), as probiotics (Deng et al., 2022; Sanders et al., 2019). A few species belonging to family *Lactobacillaceae* are detrimental microorganisms, causing spoilage (Doulgeraki et al., 2012; Gänzle, 2015; Pothakos et al., 2015; Xu et al., 2020) or behaving as opportunistic pathogens (Colautti et al., 2021; Moniente et al., 2021).

After a recent reorganization of its taxonomy (Qiao et al., 2022), the family *Lactobacillaceae* includes the vast majority of genera of lactic acid bacteria, ranging from members of the former genus *Lactobacillus*, whose species have been divided in 24 new genera (Oliphant et al., 2022; Zheng et al., 2020) and members of family *Leuconostocaceae* (*Weissella, Periweissella, Fructobacillus, Convivina, Oenococcus, Leuconostoc*). The new taxonomy is largely based on comparative genomics (Zheng et al., 2020). It allows to clearly separate genera and species on the basis of their physiology (heterofermentative vs. homofermentative species) and ecology (Duar et al., 2017; Qiao et al., 2022; Zheng et al., 2020).

The genus *Leuconostoc* has also been recently reinvestigated by comparative genomic analysis (Kumar et al., 2022) and this may lead to a reclassification of species in the near future. The ability of members of homogeneous phylogenetic groups within the family *Lactobacillaceae* to colonize related ecological niches due to specific adaptations of their physiology has been described in terms of lifestyles (Duar et al., 2017; Zheng et al., 2020). Lifestyle of *Lactobacillaceae* ranges from vertebrate-adapted (the majority of *Lactobacillus Ligilactobacillus, Limosilactobacillus*) or insect-adapted (*Apilactobacillus, Bombilactobacillus, Fructilactobacillus, Nicolia*, some *Lactobacillus*), environmental or plant associated (*Levilactobacillus*), free living (*Agrilactobacillus, Latilactobacillus, Lentilactobacillus, Levilactobacillus, Liquorilactobacillus, Paucilactobacillus, Schleiferilactobacillus, Secundilactobacillus*), nomadic (*Lacticaseibacillus, Lactiplantibacillus*) or with unknown ecology (*Companilactobacillus, Dellaglioa, Furfurilactobacillus, Holzapfelia, Lapidilactobacillus*, some *Ligilactobacillus, Loigolactobacillus*) (Duar et al., 2017; Zheng et al., 2020) (see also http://www.lactobacillus.uantwerpen.be/).

Members of the genus *Pediococcus* are frequently associated with vegetable fermentations, but can be isolated from a wide variety of habitats (Franz et al., 2014). The same applies for members of the genera *Leuconostoc* (Björkroth et al., 2014a) and *Weissella* (Björkroth et al., 2014b), while *Oenococcus* (Endo et al., 2014a), *Fructobacillus* (Endo et al., 2014b) and the new genus *Periweissella* (Bello et al., 2022) are more strictly associated with plants and vegetable fermentations.

Information on lifestyle and on history of safe use of microorganisms is important for their inclusion in lists of beneficial microorganisms and is one of the aspects EFSA considers for the inclusion (upon request) of a microbial species in the list of microorganisms with a Qualified Presumption of Safety (Bourdichon et al., 2012, 2019). The Inventory of Microbial Food Cultures (IMFC) (Bourdichon et al., 2012, 2019) has the ambition of providing a comprehensive list of microorganisms which can be used safely in food fermentations and its 2022 update (Bourdichon et al., 2022) includes 314 species, 110 of which belong to the family *Lactobacillaceae*. The criteria for inclusion in IMFC list require clear identification at the species level and demonstration, based on the scientific literature, of lack of undesirable properties (including virulence traits) and proven association of a species with a food matrix is essential.

Lifestyle and ecological niche of the former members of the genus *Lactobacillus* (Duar et al., 2017; Qiao et al., 2022; Zheng et al., 2015, 2020) have been identified using an integrated approach based on source and frequency of isolation and on occurrence of genes and metabolic traits. However, especially within the genera with a nomadic lifestyle (*Lactiplantibacillus, Lacticaseibacillus*), which colonize a large variety of animal and vegetal niches, adaptation traits have been clearly identified for different subgroups in comparative genomic studies (Douillard et al., 2013; Li et al., 2022; Wuyts et al., 2017). Further information on the lifestyle and ecology of *Lactobacillaceae* can be found in recent large metagenomic studies (Pasolli et al., 2020; Walsh et al., 2020), which also used quantitative information on prevalence and abundance to formulate and test hypotheses on lifestyle and on functional properties. Detailed prevalence and abundance data would be extremely valuable to get further insights on the distribution and specialization of *Lactobacillaceae* in foods and to identify sources of contamination.

Our research group has created and maintains the FoodMicrobionet database (Parente et al., 2016), which makes metataxonomic data on the abundance and prevalence of bacteria in foods and food related environments readily accessible. The database has been steadily growing both in size and in functions (Parente et al., 2019, 2022) and is publicly available on both GitHub (https://github.com/ep142/FoodMicrobionet/) and Mendeley data (https://data.mendeley.com/datasets/8fwwjpm79y/6).

The main objective of this work was to use the data in FoodMicrobionet to obtain quantitative insights (in terms of both abundance and prevalence) on the distribution of members of family *Lactobacillaceae* in foods and food environments. However, to do this, information on the taxonomic resolution of metataxonomic studies targeting 16S rRNA gene hypervariable regions is needed. We therefore evaluated to what extent taxonomic assignment at or below the genus level was feasible for data obtained using the most commonly used target regions within the 16S rRNA gene (V1-V3, V3-V4, V4).

## 2. Materials and methods

### 2.1 Phylogenetic analysis of 16S rRNA gene sequences of type strains of family *Lactobacillaceae*

Full length sequences of the 16S rRNA gene for type strains of members of family *Lactobacillaceae* (as defined by Zheng et al., 2020) were obtained from LPSN (https://lpsn.dsmz.de). The list of strains and sequences is shown in **Supplementary Table 1**. Whenever these sequences were too short (<1,300 bp) or had more than 5 ambiguous nucleotides, replacement sequences were obtained using GenBank or NCBI RefSeq (https://www.ncbi.nlm.nih.gov/refseq/) accession number. Sequences, as .fasta files, were imported in R (R Core Team, 2022) using functions of the package phylotools (Zhang, 2017). Shorter regions (V1-V3, V3-V4, V4) frequently used as targets in 16S metagenomics were extracted by identifying the position of commonly used primers (28F, 341F, 515F, 805R). Alignment and inference of the phylogenetic tree (for the full sequence and for regions V1-V3, V3-V4 and V4) were performed using functions of the packages DECIPHER (Wright, 2016) and phangorn (Schliep, 2011). Phylogenetic trees were annotated and plotted using functions of the packages treeio (Wang et al., 2019), tidytree (Yu, 2022), and ggtree (Yu, 2020). kmer (5-mer) distance and inference of OTUs (at 97 and 98.65% similarity) were performed using functions of the package kmer (Wilkinson, 2018).

### 2.2 Evaluation of the reliability taxonomic assignment

Taxonomy assignment was performed for full sequences and regions using functions assignTaxonomy() and addSpecies() of the dada2 package (Callahan et al., 2016) using SILVA v138.1 reference databases (https://zenodo.org/record/4587955#.YtGRqexBz0o). Confusion matrices were obtained using the function confusionMatrix() of package caret (Kuhn, 2022). In an attempt to assign reference sequences for former genus *Lactobacillus* in the SILVA database to species, kmer (5-mer) distance matrix was calculated and sequences were assigned to the species to which the type strain sequence with the lowest distance belonged. A custom reference database for assignment at the species level was built. Taxonomy assignment was performed again on both the full sequences and on V1-V3, V3-V4 and V4 regions of type strains to test the ability of the new database to obtain reliable assignments. The allowMultiple() option was used to assign, in case of ambiguous matches, up to three of the closest species matches.

### 2.3 Use of metataxonomic data to evaluate the ecological distribution of family *Lactobacillaceae* in foods and food environments

Prevalence and abundance data for all genera were extracted, tabled and plotted from FoodMicrobionet (version 4.2) for both food and food environment samples using a custom script. Only studies targeting V1-V3 or V3-V4 regions of 16S rRNA gene were used. Metadata used for annotation for the graphs included both the food sample/environment classification using the FoodEx2 system (EFSA, 2015) and data from the FoodMicrobionet samples table with information on occurrence of spoilage and fermentation.

For some genera which are represented by only one or very few species (*Dellaglioa, Holzapfelia, Schleiferilactobacillus*) further analysis of the diversity of Amplicon Sequence Variants was performed. ASVs were extracted from phyloseq objects (available at Zenodo https://doi.org/10.5281/zenodo.6954039), and further filtered on the basis of target region, length, quality etc. Annotated phylogenetic trees showing sources of ASVs and their maximum relative abundance were created as described above.

Example scripts for the bioinformatic analysis performed in this paper are available, together with supporting data files, on GitHub (https://github.com/ep142/FoodMicrobionet/tree/master/WIMB).

## 3. Results and discussion

### 3.1 Phylogenetic analysis of 16S rRNA gene sequences of type strains of family *Lactobacillaceae*

Metataxonomic studies offer great opportunities for obtaining quantitative insights in the ecology of microbial taxa in foods and other environments. The marker gene which is most frequently used for bacterial communities is the 16S rRNA gene. Due to the limitations of next generation sequencing methods, hypervariable regions (usually V1-V3, V3-V4 and V4 (Parente et al., 2022; Pollock et al., 2018) are used with amplicon length (after primer removal and merging of paired end sequences for Illumina sequencing) of <450 bp. In order to test the shortcomings of this approach in the study of the ecology of *Lactobacillaceae*, we retrieved from public databases the 16S rRNA gene sequences of type strains of family *Lactobacillaceae* and used them to infer phylogenetic trees based on full sequences and hypervariable target regions as described in Materials and Methods. The list of strains and accession number of 16S rRNA gene sequences used in this study is shown in **Supplementary Table 1**. A simplified phylogenetic tree showing the relationships between type strains of the species belonging to the family *Lactobacillaceae* is shown in **Figure 1**, while a detailed tree is shown as Supplementary Material.

**Figure 1.**
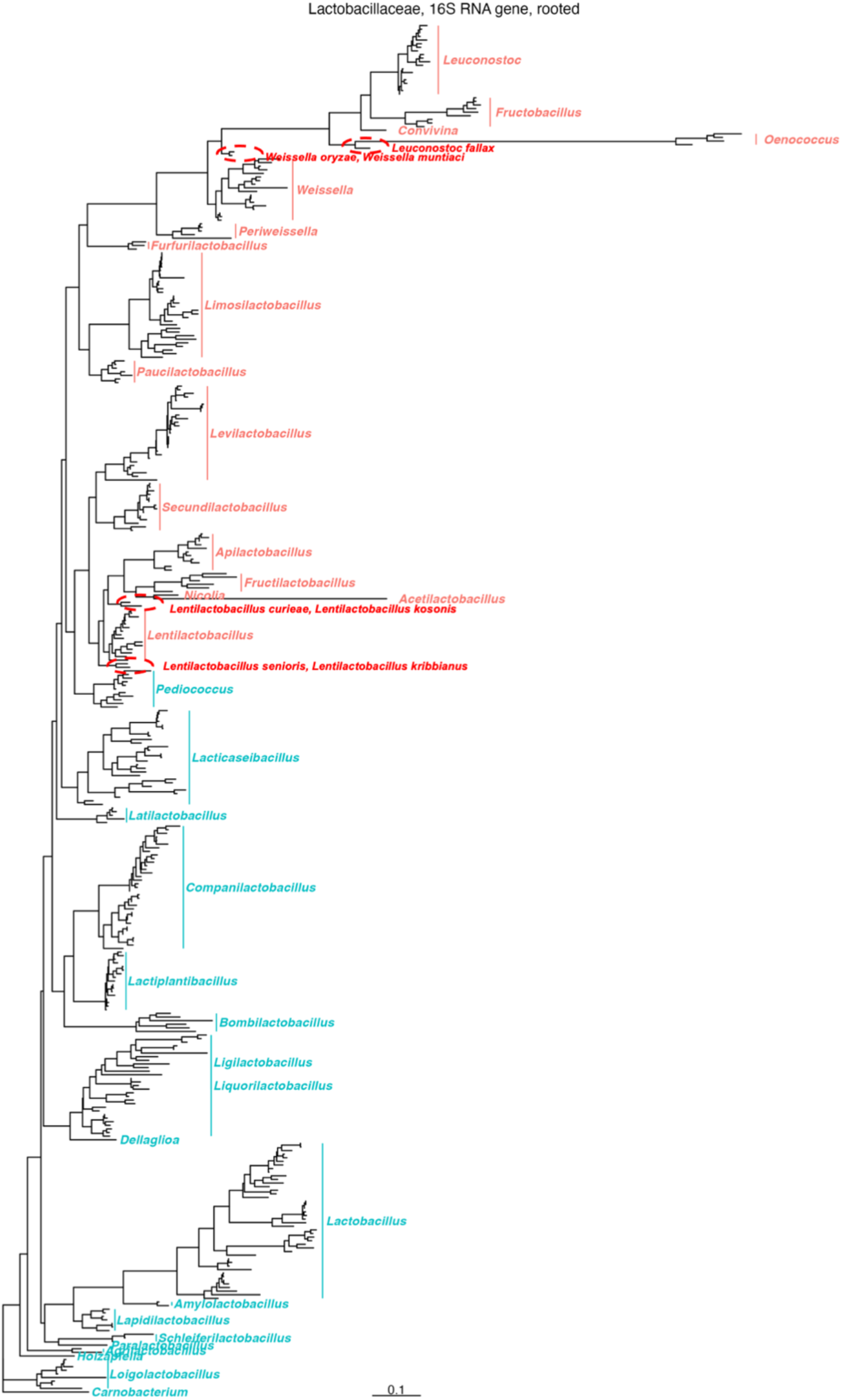
Maximum-likelihood phylogenetic tree showing the evolutionary relationships among sequences of the 16S rRNA gene for type strains of species belonging to the family *Lactobacillaceae*. Clades for each genus are outlined in blue (homofermentative species) or red (heterofermentative species) and the species which do not cluster with the corresponding genera are shown within dashed ellipses. The scale (number of nucleotide substitutions per nucleotide) is shown. Bootstrap values were 100% for all genus-level clades (data not shown).

All genera were grouped in monophyletic clusters with few exceptions (*Leuconostoc fallax, Weissella muntiaci* and *W. oryzae*, and four *Lentilactobacillus* species). As expected phylogenetic relationships differ from those inferred on the basis of WGS (Qiao et al., 2022; Zheng et al., 2020) and do not match lifestyle or metabolism. In addition, when bootstrapping was performed, bootstrapping value for genus level nodes were ≥85% with the exceptions of the nodes for genera *Lentilactobacillus, Ligilactobacillus* and *Liquorilactobacillus* (data not shown).

Use of shorter regions within the 16S rRNA results in an even worse classification, with sequences of many genera grouped in non-monophyletic clusters (**Supplementary Figures 1, 2, 3**) and is clearly worse for shorter sequences (V4). In addition, for both the V3-V4 and V4 regions many sequences were identical or almost identical.

When sequences for type strains are assigned to OTUs using kmers (5-mers), the number of OTUs retrieved at both the 97% and 98.65% (the similarity threshold at species level suggested by Qiao et al., 2022) was lower than the number of species in the family *Lactobacillaceae*. In fact, while there were 362 validly published species at the time of writing of this paper, OTUs detected using function otu() of the kmer package for the list of species in **Supplementary Table 1** at the 98.65% similarity level were only 320, 289, 211 and 174 for full length sequences, V1-V3, V3-V4 and V4 regions, respectively. In addition, for the V4 region some OTUs included species belonging to more than one genus (one OTU included *Paucilactobacillus* and *Lacticaseibacillus* and another *Agrilactobacillus, Lactiplantibacillus* and *Latilactobacillus*).

Discrimination between closely related species on the basis of full length and, to a larger extent, shorter regions of the 16S rRNA gene, was problematic with genera *Apilactobacillus, Companilactobacillus, Fructobacillus, Lacticaseibacillus, Lactiplantibacillus, Lactobacillus, Lapidilactobacillus, Latilactobacillus, Lentilactobacillus, Leuconostoc, Levilactobacillus, Liquorilactobacillus, Loigolactobacillus, Paucilactobacillus, Pediococcus, Periweissella, Secundilactobacillus, Weissella, Acetilactobacillus*.

### 3.2 Evaluation of the reliability taxonomic assignment

To obtain further information on the feasibility of taxonomic assignment at the genus and species level for full length and V1-V3, V3-V4 and V4 regions of 16S rRNA gene sequences for the family *Lactobacillaceae*, we used the SILVA v138.1 (Glöckner et al., 2017) reference database. The sensitivity and specificity of genus level assignments are shown in **Figure 2**. Data by region and genus are shown in **Supplementary Table 2**.

**Figure 2.**
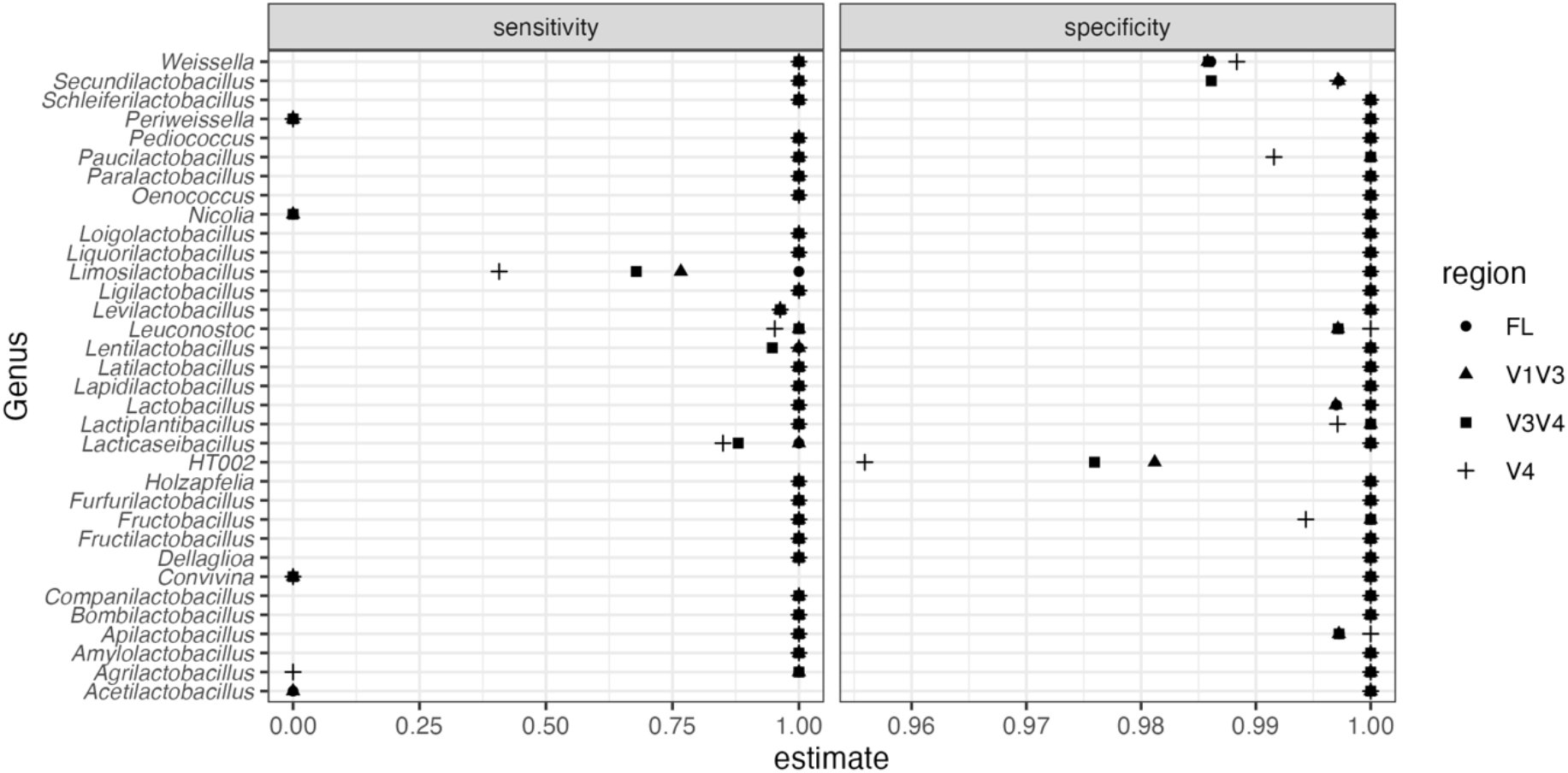
Sensitivity and specificity for taxonomic assignment of full length and V1-V3, V3V4 and V4 region sequences of the 16S rRNA gene for type strains of species belonging to family *Lactobacillaceae*. Taxonomic assignment was performed using the function assignTaxonomy() of the dada2 package using the SILVA v138.1 reference database.

The overall accuracy values for genus level assignment were: full sequence 0.975 (95% confidence interval 0.955-0.989); V1-V3 0.957 (0.931-0.975), V3-V4 0.944 (0.915-0.964), V4 0.926 (0.893-0.950). Accuracy clearly decreased as the length of the target sequence decreased and, for comparable lengths, was worse for the V3-V4 region compared to the V1-V3 region. The 0 value for sensitivity for genera *Acetilactobacillus, Convivina, Nicolia* and *Periweissella* reflects the lack of sequences assigned to these genera in the SILVA v138.1 database. Low level of sensitivity for shorter sequences of genus *Limosilactobacillus* and low levels of specificity for sequences labelled as “HT002” in the SILVA database were due to taxonomic assignment of sequences of some type strains of *Limosilactobacillus* to “HT002”. Relatively low values for specificity in taxonomic assignment of short sequences for *Secundilactobacillus* was due to the erroneous identification of some sequence of type strains of *Levilactobacillus, Lacticaseibacillus*, and *Lentilactobacillus* as belonging to *Secundilactobacillus*. Low specificity for *Weissella* was systematically due to the species which now belong to genus *Periweissella*.

We conclude that, with few exceptions, taxonomic assignment at the genus level of 16S rRNA genes sequences using the naïve Bayes classifier and the SILVA v138.1 reference database can be done for almost all genera of the family *Lactobacillaceae* with a very high degree of confidence for full length sequences, and with a satisfactory level of accuracy for the V1-V3 regions. Results for the V3-V4 and V4 regions are still acceptable but significantly worse. This confirms the results of Bukin et al. (2019).

Lifestyle appears to be relatively constant within genera of the family *Lactobacillaceae* (Duar et al., 2017; Qiao et al., 2022; Zheng et al., 2020). On the other hand, several important genera have been found in a wide variety of niches, and, although their lifestyle has been defined as nomadic, evidence of niche adaptation exists for well-defined clades within important species (Douillard et al., 2013; Li et al., 2022; Wuyts et al., 2017). If species assignment was possible with a good degree of confidence in metataxonomic studies, one would be able to exploit databases such as FoodMicrobionet to get a deeper insight on the distribution of individual species within the family *Lactobacillaceae*.

Since the SILVA v138.1 database does not allow species assignment for sequences belonging to the former genus *Lactobacillus* we created a custom database in which the reference sequences for the 23 genera proposed by Zheng et al. (2020) were attributed to the same species of the type strain sequence with the smallest kmer distance. We then performed taxonomic assignment at the genus and species level for type strain sequences using functions of package dada2 (Callahan et al., 2016). The allowMultiple option was used to obtain multiple matches whenever a single, unambiguous match was not possible. The proportion of matches, partial matches and failure to match species identification is shown in **Figure 3**.

**Figure 3.**
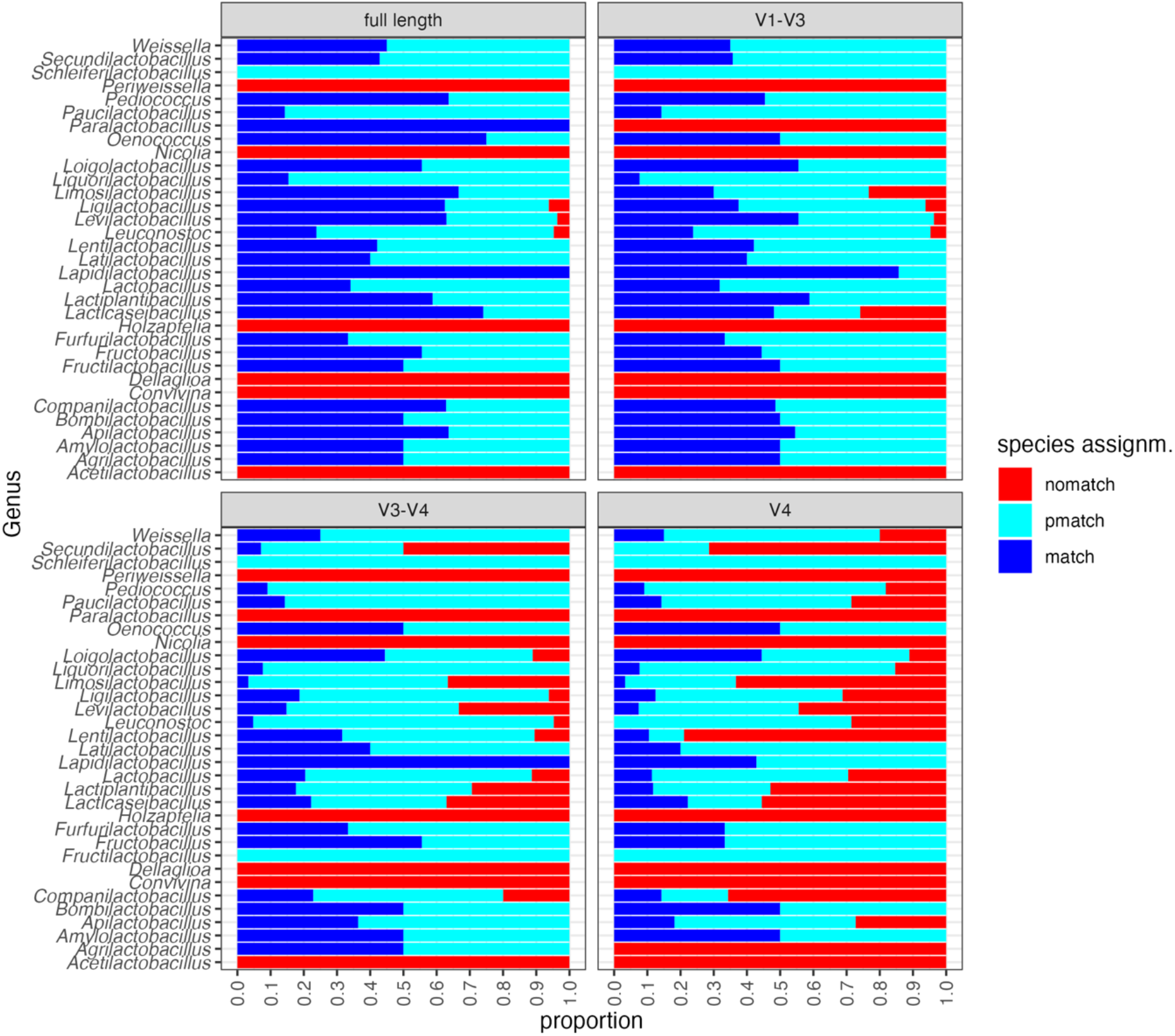
Proportion of correct taxonomic assignments for full-length and partial sequences (V1-V3, V3-V4, V4) of the 16S rRNA gene for type strains of the family *Lactobacillaceae*. Species assignment was performed using the addSpecies() function of the dada2 package, using a custom reference database derived from SILVA v138.1. match: the species assignment was unambiguous; pmatch the species assignment was ambiguous (more than one match returned by the naïve Bayes classifier) but included the correct identification; nomatch: no species assignment was returned.

The distribution within-genus of the proportion of matching species identifications is shown in **Supplementary Figure 4**. The median value for the within-genus proportion of matching species identification was 0.500, 0.375, 0.176 and 0.105 for full length sequences, V1-V3, V3V4 and V4, respectively.

We conclude that taxonomic assignment down to the species level using the naïve Bayes classifier for sequences for the V1-V3, V3-V4, V4 regions of the 16S rRNA gene of members of the family *Lactobacillaceae* is hardly possible, and that, even for full length sequences, only 50% of the type strain sequences can be unambiguously assigned to species. This low percentage of successful identification may be due to the relatively large number of closely related species in some genera *(Lactiplantibacillus, Lacticaseibacillus*, etc.) for which the within-genus similarity of 16S rRNA gene sequences is very high.

This confirms that use of 16S rRNA gene targets in metataxonomic studies can reliably resolve genus, but not species, unless the full-length gene is used (Bokulich et al., 2020; Johnson et al., 2019; Pollock et al., 2018). Custom reference databases have been claimed to improve the accuracy of taxonomic assignment (Meola et al., 2019) and, recently, the *cpn60* gene has demonstrated potential in the identification at the species level of short sequences of *Lactobacillaceae* (Links et al., 2012; Shukla and Hill, 2022). Alternatives to the naïve Bayes classifier (Wang et al., 2007) have also been claimed to improve taxonomic assignment when using targeted metagenomics (Murali et al., 2018; Ziemski et al., 2021). Further testing is needed to identify the best tool for the identification of *Lactobacillaceae* at the species level using metataxonomic data.

### 3.3 Use of metataxonomic data to evaluate the ecological distribution of family *Lactobacillaceae* in foods and food environments

The vast majority of metataxonomic studies still use short sequences of the 16S rRNA gene, and these data, when properly organized, can be exploited to evaluate the distribution of genera of *Lactobacillaceae* in foods and food environments.

We therefore exploited FoodMicrobionet for this purpose. *Lactobacillaceae* are found in 194 studies out of 211 and in 8,007 food or food environment samples out of 12,748. This confirms that they are a ubiquitous group. A prevalence and abundance table is provided in Supplementary Material.

Further analysis was carried out separately for food samples and food environment samples. In a first step, we limited the analysis to food samples, with longer sequences (>300 bp) targeting the V1-V3 and V3-V4 regions, 125 studies and 5,393 samples and calculated prevalence and abundance for the L1 category of the EFSA FoodEx2 classification (EFSA, 2015, see **Supplementary Table 3**), which groups foods in large categories. This is a very conservative choice, because, for many genera, even shorter sequences (V4 region) would guarantee a reliable genus level identification. The results are shown in **Figure 4**.

**Figure 4.**
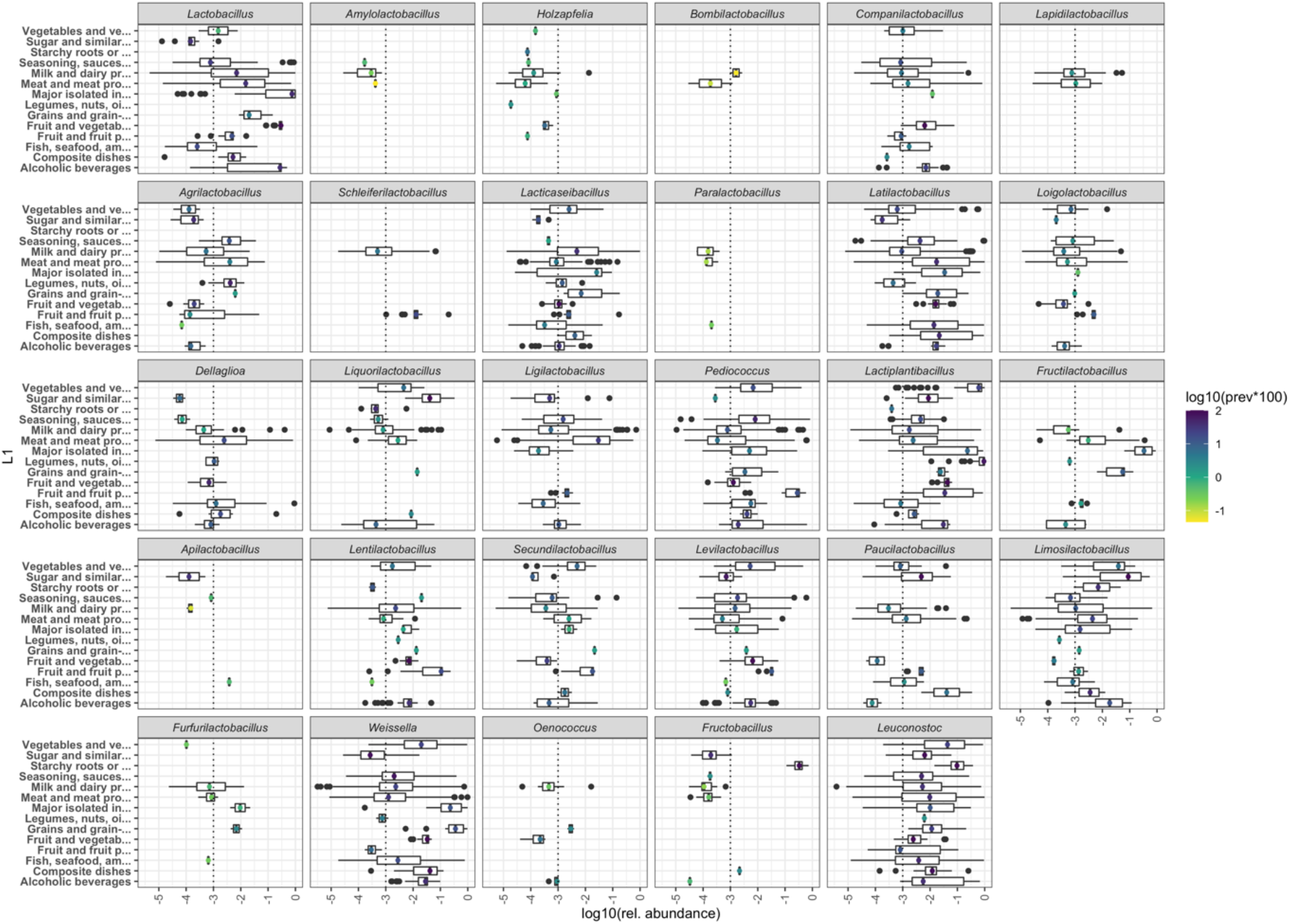
Boxplot showing the distribution of abundance of sequences (V1-V3 and V3-V4, length >350 bp) of genera of the family *Lactobacillaceae* in food samples. The prevalence is shown as the colour scale of a marker centred on the median. Foods are grouped on the basis of the L1 category of the EFSA FoodEx2 classification (EFSA, 2015). Names of the categories have been shortened in some cases. Refer to **Supplementary Table 3**. Genera in facets were ordered on the basis of their taxonomic relationships (Qiao et al., 2022; Zheng et al., 2020).

The level of aggregation provided by L1 may be too coarse for understanding the ecology of individual genera. We therefore recalculated prevalence and statistics for abundance for the L4 level of the FoodEx 2 classification. Obtaining a plot for all genera using this aggregation level is prohibitive because there are too many categories. However, a prevalence and abundance table is provided as Supplementary Material. Example plots for a few generalist (*Lacticaseibacillus, Latilactobacillus, Dellaglioa, Levilactobacillus, Weissella*) and specialist (*Holzapfelia, Schleiferilactobacillus, Liquorilactobacillus, Fructilactobacillus, Fructobacillus*) genera are shown in **Supplementary Figures 5** to **14**.

A few genera of the family *Lactobacillaceae* rarely appear in the sample set after filtering and almost never exceed a relative abundance of 0.01. These include *Amylolactobacillus* (unknown lifestyle in Zheng et al., 2020), *Holzapfelia* (unknown lifestyle in Zheng et al., 2020, but found with low prevalence and abundance in bovine muscle, cattle milk and cheese), *Bombilactobacillus* (insect associated), *Paralactobacillus* (unknown lifestyle), *Apilactobacillus* (insect associated). Since the original source of isolation of *Holzapfelia floricola* (the only species of this genus) was a flower and since *Bombilactobacillus* is insect associated, their presence in bovine meat or milk samples may be associated with pasture.

A few genera were clearly associated with specific food environments, in which they occasionally appeared with relatively high abundance. *Lapidilactobacillus* was found in dairy and meat products, although its lifestyle was described as unknown by Zheng et al. (2020). Type strains of this genus were isolated from pickles, distilled spirits, silage (as found in the BacDive database https://bacdive.dsmz.de/). Its origin may therefore be vegetable sources. *Schleiferilactobacillus* occurred in milk and fermented milk products, with occasionally high relative abundance (0.1-0.2) in firm ripened cheeses which were also spoiled and in fermented fruit products (**Supplementary Figure 7**). Its lifestyle has been described as free-living and type strains have been isolated from fermented vegetable products. *Dellaglioa* sequences were retrieved sometimes with a high prevalence and abundance from a variety of sources, and their relative abundance was highest in meat and meat products, with high abundances often associated with spoiled products (**Supplementary Figure 10**). The only species of this genus is *Dellaglioa algida*, which has an unknown lifestyle but whose strains, in accordance with our findings, have been isolated from vacuum packaged meat. This genus may therefore be animal associated. *Fructilactobacillus* sequences were found in a variety of food categories, with highest abundances in yeast cultures and sourdoughs, which is compatible with the ecology of *Fructilactobacillus sanfranciscensis* (De Vuyst et al., 2014). This genus has been described as insect associated (Zheng et al., 2020). The significance of the occurrence, at low abundance, in milk, meat and smoked fish products is unclear and may due to contaminating sequences. *Furfurilactobacillus* is another genus whose sequences have a restricted distribution. Its lifestyle is unknown, although the source of type strains is invariably sourdough. In agreement with this, we found that its highest median abundance (which was close to 0.01) was found in flours and sourdoughs. Again, occurrence of sequences in products of animal origin has an unclear significance. *Oenococcus* is a genus which is strictly adapted to alcoholic beverages (Badotti et al., 2014; Bech-Terkilsen et al., 2020; Endo and Okada, 2006); the occurrence of sequences assigned to this genus in foods of animal origin may be certainly suspicious. Members of the genus *Fructobacillus* are fructophilic LAB which are associated with fructose-rich environments (Dicks and Endo, 2022; Endo and Salminen, 2013; Endo et al., 2018) including fruits and honeybees and honey. The abundance of their sequences in vegetable products, with high abundance in sugar plants is therefore not surprising, while their occasional abundance in milk and dairy products is unclear.

Many other genera are definitely generalist, and their sequences occur with sometimes high prevalence and abundance in a large variety of foods of plant and animal origin (**Figure 4**). These include *Lactobacillus* (lifestyle described as vertebrate, animal or insect associated), a genus including several species used as starters in dairy fermentations (Bourdichon et al., 2022), *Companilactobacillus, Lacticaseibacillus, Latilactobacillus, Ligilactobacillus, Pediococcus, Lactiplantibacillus, Lentilactobacillus, Secundilactobacillus, Levilactobacillus, Paucilactobacillus, Leuconostoc, Weissella*. This is in good agreement with their nomadic or free-living lifestyles. *Loigolactobacillus* and *Limosilactobacillus* have been described as vertebrate associated, but sequences assigned to these genera are widely distributed in food samples, including those of vegetable origin.

The sources of *Lactobacillaceae* in foods can be many and diverse. Although source tracking has been used for a few studies, and environmental microbiomes, especially in traditional food productions, are colonized by stable communities which shape the microbial landscape of different areas of food factories (da Silva Vale et al., 2021; De Filippis et al., 2020; Sun and D’Amico, 2021), unambiguous attribution of sources of contamination can be difficult due to the low taxonomic resolution of metataxonomic studies. As a proof of concept, we performed a search in the FoodMicrobionet database using the same filtering criteria described above. Only 19 studies with 908 samples matched these criteria and the results are shown in **Figure 5**. The number of food environment samples in FoodMicrobionet is much lower than that of food samples (only 2,197 out of 12,145) and the coverage of food categories is far less homogeneous. In addition, the description of food environmental samples is not standardized. Here, we have tried to extract some common naming patterns, but the number of samples is too low to create further subcategories: although “teat” is always referred to dairy cow’s teat surfaces, all the other items may refer to environments related to different food industries or raw material production contexts. Anyway, while it is quite clear that generalist like *Lactobacillus, Lacticaseibacillus, Latilactobacillus, Lactiplantibacillus, Weissella* and *Leuconostoc* are widely distributed in food environments and sometime are abundant components (>0.1 relative abundance) of the microbiota, the number of variety of samples is too low to evaluate associations of genera with potential sources of contamination.

**Figure 5.**
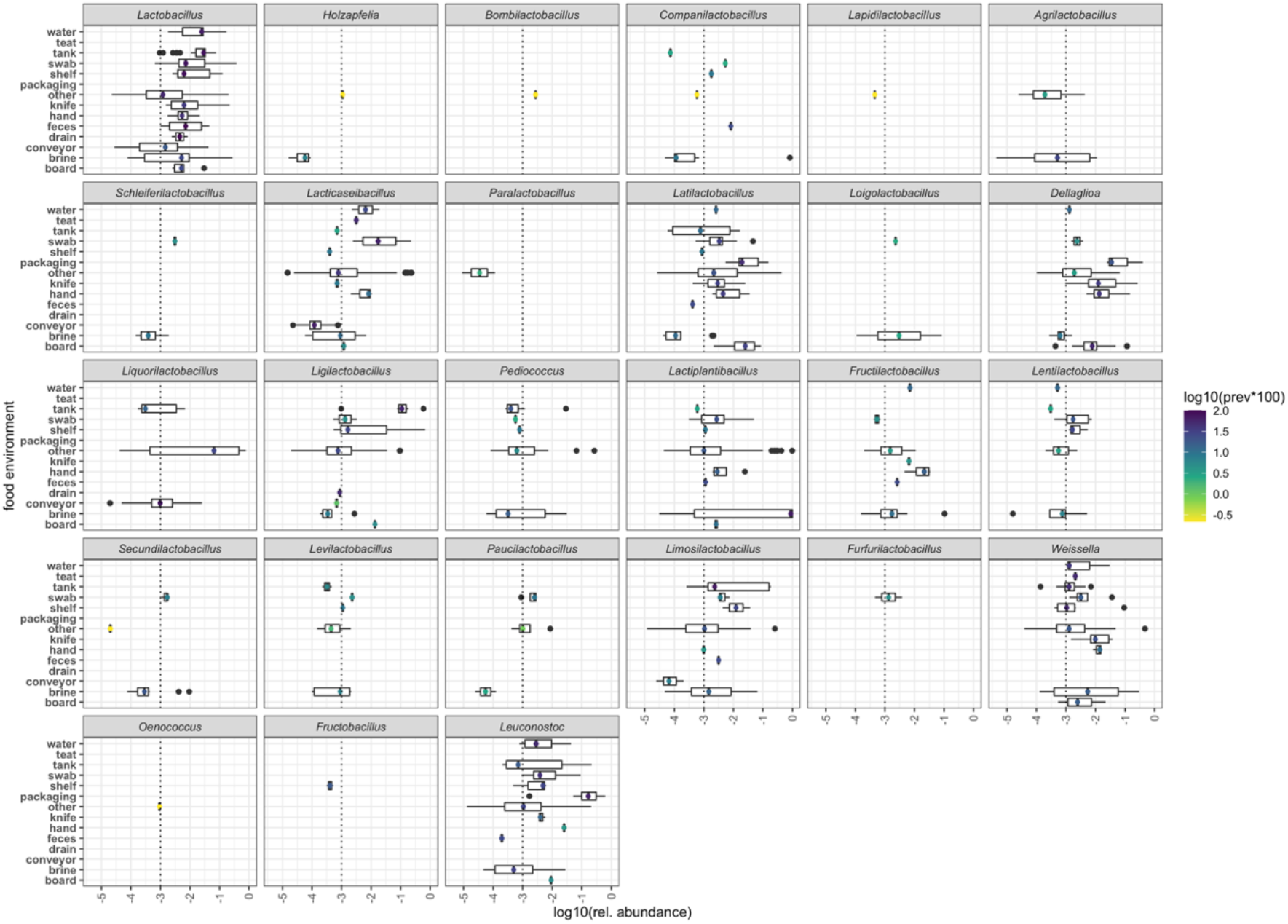
Boxplot showing the distribution of abundance of sequences (V1-V3 and V3-V4, length >350 bp) of genera of the family *Lactobacillaceae* in food related environmental samples. The prevalence is shown as the colour scale of a marker centred on the median. Genera in facets were ordered on the basis of their taxonomic relationships (Qiao et al., 2022; Zheng et al., 2020).

Starting from version 4.1, FoodMicrobionet includes, as an external resource (https://doi.org/10.5281/zenodo.6954040), R lists containing the ASV for all studies processed using the dada2 pipeline (Parente et al., 2022). Although the ecological significance of ASV is not clear (i.e., there is no convincing proof that they represent different subpopulations within a taxonomic group) they have been used to evaluate microdiversity in some studies (Quijada et al., 2018; Skeie et al., 2019; Zwirzitz et al., 2020). Their main advantage, however, is that results from different studies can be compared, something which is impossible when using Operational Taxonomic Units (Callahan et al., 2017). We used ASV analysis to evaluate the distribution of ASVs for the three genera of family *Lactobacillaceae* which have only one or very few species and which are relatively prevalent or abundant in at least some food groups (*Dellaglioa, Holzapfelia, Schleiferilactobacillus*). The results are shown in **Supplementary Figures 15** to **20**. The results confirm the findings obtained in previous sections. Only occasionally ASV did not cluster with the corresponding reference sequences, thus confirming the reliability of the taxonomic assignment. On the other hand, the diversity of partial sequences was usually very low and discriminating ASVs on the basis of their clustering with the type strains of a given species is difficult if not impossible (see for example **Supplementary Figures 19** and **20**, for *Schleiferilactobacillus*). In one instance (**Supplementary Figure 18**, *Holzapfelia*, V3-V4) a number of ASVs provided a hint of belonging to a different, possibly unknown species, but the supporting evidence is too scant. In some instances, the grouping of ASVs was clearly confounded with the study from which they were obtained. No clear evidence of association of sub-clusters of ASVs with a given food source was obtained.

## 4. Conclusions

In this paper we have demonstrated that the FoodMicrobionet metataxonomic database can be used to retrieve detailed quantitative information on the distribution of economically important taxa (here genera belonging to the family *Lactobacillaceae*). In a preliminary analysis we confirmed that taxonomic assignment at the genus level was effective (with a few exceptions) using full length sequences of the 16S rRNA gene and to a lesser extent, with partial sequences (with longer sequences, V1-V3 and V3-V4, performing much better than V4), while assignment at the species level with short sequences was rarely possible. This information may guide researchers in the choice of the target regions in metataxonomic studies for foods or environments in which *Lactobacillaceae* are important components of the microbiota. Use of short regions, like V4, should altogether be avoided if genera for which specificity and sensitivity of taxonomic assignment is low are considered to be important.

We therefore used a very conservative approach, based on the analysis of samples for which sequencing was carried out using longer target regions (V1-V3 and V3-V4), for investigating the ecological distribution of family *Lactobacillaceae* in foods. Our results provide support to the description of lifestyles of *Lactobacillaceae* (Duar et al., 2017; Zheng et al., 2020) and provides hints on the ecological niches of rarely isolated genera. The approach can be easily replicated for other taxonomic groups.

The biological significance of conclusion based on metataxonomic approaches targeting DNA should be evaluated with caution. It is well known that the active fraction of the food microbiota may be significantly different from the total microbiota, whose composition is evaluated using DNA as a target (Erkus et al., 2013, 2016) and that contamination during wet laboratory procedures and traces of DNA from ingredients may influence the composition of the microbiota (Dahlberg et al., 2019; Duthoo et al., 2022). Although recent studies sometimes include reagent blanks, the information in sample metadata tables obtained from NCBI Short Read Archive is not sufficient for statistical removal of contaminating sequences (Davis et al., 2018). One conservative approach to address the second problem might be to remove samples in which the relative abundance of a target taxon is very low (<10^−3^). However, this may significantly reduce the sensitivity of the analysis and arbitrarily remove rare taxa. Finally, we found that combining ASVs from different studies (Callahan et al., 2017) is indeed possible, but provides little additional information, even when strict criteria are used for the filtering of sequences.

## Supporting information

Prevalence and abundance data

Prevalence and abundance data L4 level FoodEx2

xlsx version of supplementary table 1

## Acknowledgements

This work was funded by the European Commission – NextGenerationEU, Project “Strengthening the MIRRI Italian Research Infrastructure for Sustainable Bioscience and Bioeconomy”, code n. “IR0000005 (within the Italy’s National Recovery and Resilience Plan).

---

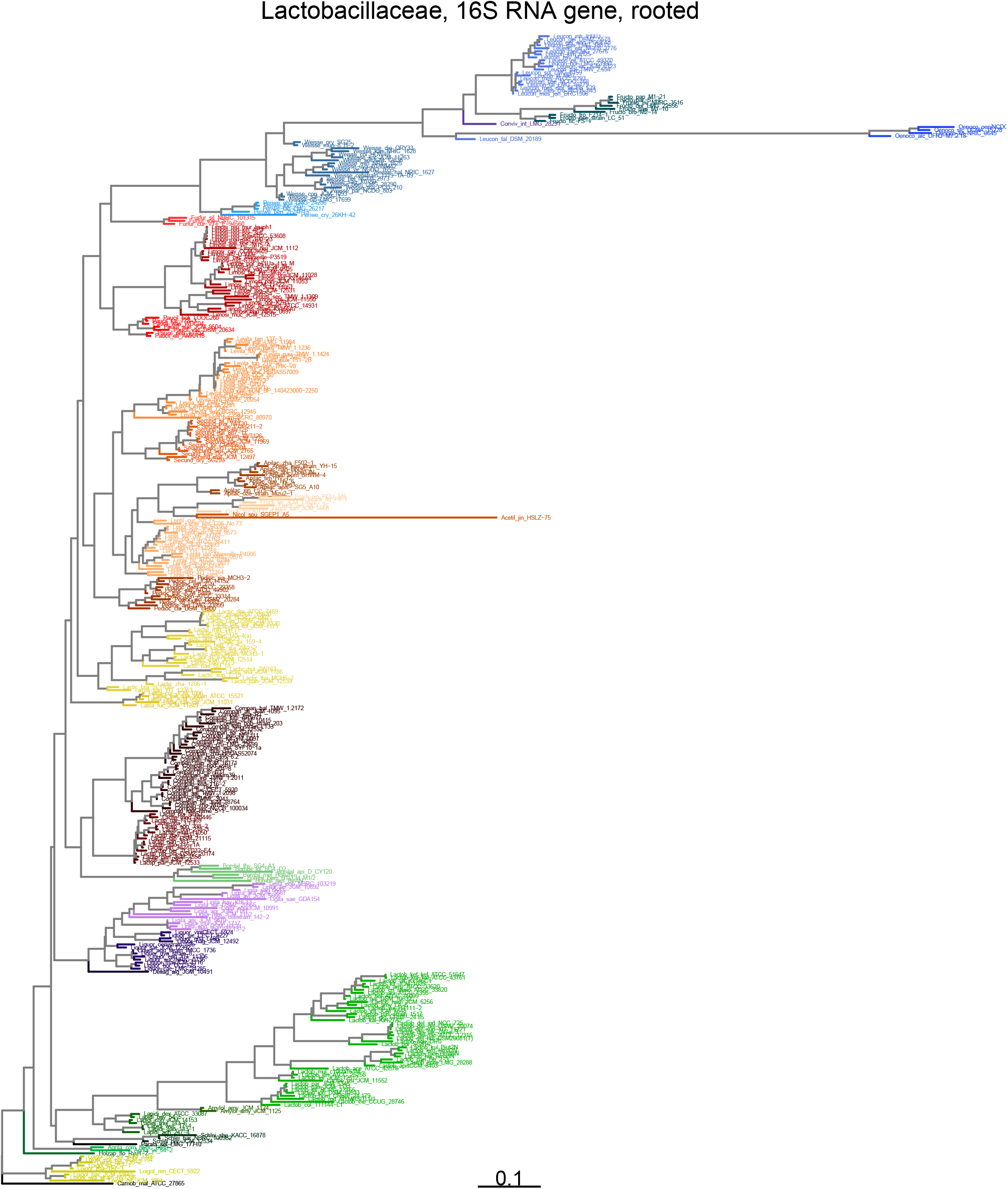

