## Supplementary tables and figures for "Metataxonomic insights in the distribution of *Lactobacillaceae* in foods and food environments"

Scuola di Scienze Agrarie, Forestali, Alimentari ed Ambientali, Università degli Studi della Basilicata, Potenza, Italy

*Corresponding author
Prof. Eugenio Parente


### Supplementary Tables and Figures

**Supplementary Table 1**. List of species and type strains and 16S rRNA gene sequence accession numbers of family *Lactobacillaceae* used in this study. The table is a compilation of information from http://www.lactobacillus.uantwerpen.be/ and from https://lpsn.dsmz.de/. *Carnobacterium maltaromaticum* was included as an outgroup.

| Genus | Species | Subspecies | Strain | Seq. accession[1] | Lifestyle | Ferm. type[2] | Abbr.[3] |
|---|---|---|---|---|---|---|---|
| *Acetilactobacillus* | *jinshanensis* | | HSLZ-75 | KT783533 | Unknown | HE | Acetil_jin_HSLZ-75 |
| *Agrilactobacillus* | *composti* | | NRIC_0689 | AB268118 | Free Living | HO | Agrila_com_NRIC_0689 |
| *Agrilactobacillus* | *yilanensis* | | 54-2 | MK110806 | Free Living | HO | Agrila_yil_54-2 |
| *Amylolactobacillus* | *amylophilus* | | JCM_1125 | NR_044702.2 | Unknown | HO | Amylol_amy_JCM_1125 |
| *Amylolactobacillus* | *amylotrophicus* | | JCM_1124 | NR_042511.1 | Unknown | HO | Amylol_amy_JCM_1124 |
| *Apilactobacillus* | *apinorum* | | Fhon13N | JX099541 | Insect | HE | Apilac_api_Fhon13N |
| *Apilactobacillus* | *apisilvae* | | SG5_A10 | OM986478 | Insect | HE | Apilac_apis__SG5_A10 |
| *Apilactobacillus* | *bombintestini* | | BHWM-4 | CP032626 | | HE | Apilac_bom_BHWM-4 |
| *Apilactobacillus* | *kunkeei* | | strain_YH-15 | Y11374 | Insect | HE | Apilac_kun_strain_YH-15 |
| *Apilactobacillus* | *micheneri* | | Hlig3 | KT833121 | Insect | HE | Apilac_mic_Hlig3 |
| *Apilactobacillus* | *nanyangensis* | | HN36-1 | LC597577 | | HE | Apilac_nan_HN36-1 |
| *Apilactobacillus* | *ozensis* | | strain_Mizu2-1 | AB572588 | Insect | HE | Apilac_oze_strain_Mizu2-1 |
| *Apilactobacillus* | *quenuiae* | | HV_6 | KX656667 | Insect | HE | Apilac_que_HV_6 |
| *Apilactobacillus* | *timberlakei* | | HV_12 | KX656650 | Insect | HE | Apilac_tim_HV_12 |
| *Apilactobacillus* | *xinyiensis* | | F575-4 | LC634436 | Insect | HE | Aplilac_xin_F575-4 |

[1] Accession number for the 16S rRNA gene sequence used in this study
[2] HE heterofermentative, HO homofermentative
[3] abbreviation used in dendrograms

| | | | | | | |
|---|---|---|---|---|---|---|
| *Apilactobacillus* | *zhangqiuensis* | F502-1 | LC634435 | Insect | HE | Apilac_zha_F502-1 |
| *Bombilactobacillus* | *apium* | D_CY120 | MN709051 | | HE | Bombil_api_D_CY120 |
| *Bombilactobacillus* | *bombi* | BTLCH_M1/2 | KJ078643 | Insect | HO | Bombil_bom_BTLCH_M1/2 |
| *Bombilactobacillus* | *folatiphilus* | SG4_D2 | OM986477 | Insect | HO | Bombil_fol_SG4_D2 |
| *Bombilactobacillus* | *mellifer* | Bin4N | JX099543 | Insect | HO | Bombil_mel_Bin4N |
| *Bombilactobacillus* | *mellis* | Hon2N | JX099545 | Insect | HO | Bombil_mel_Hon2N |
| *Bombilactobacillus* | *thymidiniphilus* | SG4_A1 | OM986476 | Insect | HO | Bombil_thy_SG4_A1 |
| *Companilactobacillus* | *alimentarius* | JCM_1095 | NR_044701.2 | Unknown | HO | Compan_ali_JCM_1095 |
| *Companilactobacillus* | *allii* | WiKim39 | KY419164 | Unknown | HO | Compan_all_WiKim39 |
| *Companilactobacillus* | *baiquanensis* | 184-8 | MK110828 | Unknown | HO | Compan_bai_184-8 |
| *Companilactobacillus* | *bobalius* | strain_203 | AY681134 | Unknown | HO | Compan_bob_strain_203 |
| *Companilactobacillus* | *crustorum* | LMG_23699 | AM285450 | Unknown | HO | Compan_cru_LMG_23699 |
| *Companilactobacillus* | *farciminis* | JCM_1097 | NR_044707.2 | Unknown | HO | Compan_far_JCM_1097 |
| *Companilactobacillus* | *formosensis* | S215 | AB794060 | Unknown | HO | Compan_for_S215 |
| *Companilactobacillus* | *furfuricola* | JCM_18764 | AB910349 | Unknown | HO | Compan_fur_JCM_18764 |
| *Companilactobacillus* | *futsaii* | YM_0097 | HQ322270 | Unknown | HO | Compan_fut_YM_0097 |
| *Companilactobacillus* | *ginsenosidimutans* | EMML_3041 | HQ389549 | Unknown | HO | Compan_gin_EMML_3041 |
| *Companilactobacillus* | *halodurans* | TMW_1.2172 | MK968448 | Unknown | HO | Compan_hal_TMW_1.2172 |

| | | | | | | |
|---|---|---|---|---|---|---|
| *Companilactobacillus* | *heilongjiangensis* | S4-3 | JF411966 | Unknown | HO | Compan_hei_S4-3 |
| *Companilactobacillus* | *huachuanensis* | 395-6.2 | LC438522 | Unknown | HO | Compan_hua_395-6.2 |
| *Companilactobacillus* | *hulinensis* | 8-1(1) | MK110830 | Unknown | HO | Compan_hul_8-1(1) |
| *Companilactobacillus* | *insicii* | TMW_1.2011 | KP677494 | Unknown | HO | Compan_ins_TMW_1.2011 |
| *Companilactobacillus* | *jidongensis* | 204-8 | MK110829 | Unknown | HO | Compan_jid_204-8 |
| *Companilactobacillus* | *kedongensis* | 116-2 | MK110817 | Unknown | HO | Compan_ked_116-2 |
| *Companilactobacillus* | *keshanensis* | 33-7 | MK110816 | Unknown | HO | Compan_kes_33-7 |
| *Companilactobacillus* | *kimchiensis* | strain_L133 | HQ906500 | Unknown | HO | Compan_kim_strain_L133 |
| *Companilactobacillus* | *kimchii* | AP1077 | AF183558 | Unknown | HO | Compan_kim_AP1077 |
| *Companilactobacillus* | *metriopterae* | Hime_5-1 | LC190736 | Unknown | HO | Compan_met_Hime_5-1 |
| *Companilactobacillus* | *mindensis* | JCM_12532 | NR_028949.1 | Unknown | HO | Compan_min_JCM_12532 |
| *Companilactobacillus* | *mishanensis* | 256-3 | MK110831 | Unknown | HO | Compan_mis_256-3 |
| *Companilactobacillus* | *musae* | 313 | LC184607 | Unknown | HO | Compan_mus_313 |
| *Companilactobacillus* | *nantensis* | JCM_16171 | LC258154 | Unknown | HO | Compan_nan_JCM_16171 |
| *Companilactobacillus* | *nodensis* | iz4b-1 | AB332024 | Unknown | HO | Compan_nod_iz4b-1 |
| *Companilactobacillus* | *nuruki* | SYF10-1a | KY711184 | Unknown | HO | Compan_nur_SYF10-1a |
| *Companilactobacillus* | *pabuli* | NFFJ11 | MT114679 | | HO | Compan_pab_NFFJ11 |
| *Companilactobacillus* | *paralimentarius* | JCM_10415 | NR_114844.1 | Unknown | HO | Compan_par_JCM_10415 |
| *Companilactobacillus* | *salsicarnum* | TMW_1.2098 | MK968446 | Unknown | HO | Compan_sal_TMW_ |

| | | | | | | |
|---|---|---|---|---|---|---|
| | | | | | | 1.2098 |
| *Companilactobacillus* | *suantsaicola* | R7 | MH810311 | Unknown | HO | Compan_sua_R7 |
| *Companilactobacillus* | *tucceti* | CECT_5920 | AJ576006 | Unknown | HO | Compan_tuc_CECT_5920 |
| *Companilactobacillus* | *versmoldensis* | NCCB_100034 | NR_028990.1 | Unknown | HO | Compan_ver_NCCB_100034 |
| *Companilactobacillus* | *zhachilii* | HBUAS52074 | MH392835 | Unknown | HO | Compan_zha_HBUAS52074 |
| *Companilactobacillus* | *zhongbaensis* | M1575 | MK110832 | Unknown | HO | Compan_zho_M1575 |
| *Convivina* | *intestini* | LMG_28291 | LK054488 | | HE | Conviv_int_LMG_28291 |
| *Dellaglioa* | *algida* | JCM_10491 | NR_028617.1 | Unknown | HO | Dellag_alg_JCM_10491 |
| *Fructilactobacillus* | *florum* | F9-1 | AB498045 | Insect | HE | Fructi_flo_F9-1 |
| *Fructilactobacillus* | *fructivorans* | ATCC_8288 | NR_036789.1 | Insect | HE | Fructi_fru_ATCC_8288 |
| *Fructilactobacillus* | *ixorae* | PCU_346 | MT758158 | Insect | HE | Fructi_ixo_PCU_346 |
| *Fructilactobacillus* | *lindneri* | JCM_11027 | NR_029308.2 | Insect | HE | Fructi_lin_JCM_11027 |
| *Fructilactobacillus* | *sanfranciscensis* | JCM_5668 | NR_029261.2 | Insect | HE | Fructi_san_JCM_5668 |
| *Fructilactobacillus* | *vespulae* | DCY_75 | JX863367 | Insect | HE | Fructi_ves_DCY_75 |
| *Fructobacillus* | *broussonetiae* | M2-14 | MT026984 | | HE | Fructo_bro_M2-14 |
| *Fructobacillus* | *durionis* | LMG_22556 | AJ780981 | | HE | Fructo_dur_LMG_22556 |
| *Fructobacillus* | *ficulneus* | FS-1 | AF360736 | | HE | Fructo_fic_FS-1 |
| *Fructobacillus* | *fructosus* | NBRC_3516 | AB680098 | | HE | Fructo_fru_NBRC_3516 |
| *Fructobacillus* | *papyriferae* | M1-10 | MT026981 | | HE | Fructo_pap_M1-10 |

| | | | | | | |
|---|---|---|---|---|---|---|
| *Fructobacillus* | *papyrifericola* | M1-21 | MT026983 | | HE | Fructo_pap_M1-21 |
| *Fructobacillus* | *parabroussonetiae* | S1-1 | MT026985 | | HE | Fructo_par_S1-1 |
| *Fructobacillus* | *pseudoficulneus* | strain_LC_51 | AY169967 | | HE | Fructo_pse_strain_LC_51 |
| *Fructobacillus* | *tropaeoli* | F214-1 | AB542054 | | HE | Fructo_tro_F214-1 |
| *Furfurilactobacillus* | *curtus* | VTT_E-94560 | LC093898 | Unknown | HE | Furfur_cur_VTT_E-94560 |
| *Furfurilactobacillus* | *rossiae* | CS1 | AJ564009 | Unknown | HE | Furfur_ros_CS1 |
| *Furfurilactobacillus* | *siliginis* | NBRC_101315 | AB370882 | Unknown | HE | Furfur_sil_NBRC_101315 |
| *Holzapfelia* | *floricola* | Ryu1-2 | AB523780 | Unknown | HO | Holzap_flo_Ryu1-2 |
| *Lacticaseibacillus* | *absianus* | YH-lac23 | MT009028 | | HO | Lactic_abs_YH-lac23 |
| *Lacticaseibacillus* | *baoqingensis* | 47-3 | MK110840 | Unknown | HO | Lactic_bao_47-3 |
| *Lacticaseibacillus* | *brantae* | SL1108 | HQ022861 | Unknown | HO | Lactic_bra_SL1108 |
| *Lacticaseibacillus* | *camelliae* | strain_MCH3-1 | AB257864 | Unknown | HO | Lactic_cam_strain_MCH3-1 |
| *Lacticaseibacillus* | *casei* | DSMZ_20011 | NR_041893.1 | Nomadic | HO | Lactic_cas_DSMZ_20011 |
| *Lacticaseibacillus* | *chiayiensis* | BCRC_81062 | MF446960 | Unknown | HO | Lactic_chi_BCRC_81062 |
| *Lacticaseibacillus* | *daqingensis* | 143-4(a) | MK110842 | Unknown | HO | Lactic_daq_143-4(a) |
| *Lacticaseibacillus* | *hegangensis* | 73-4 | MK110833 | Unknown | HO | Lactic_heg_73-4 |
| *Lacticaseibacillus* | *hulanensis* | ZW163 | LC436604 | Free Living | HO | Lactic_hul_ZW163 |
| *Lacticaseibacillus* | *jixianensis* | 159-4 | MK110836 | Unknown | HO | Lactic_jix_159-4 |
| *Lacticaseibacillus* | *manihotivorans* | JCM_12514 | NR_024835.1 | Unknown | HO | Lactic_man_JCM_12514 |
| *Lacticaseibacillus* | *mingshuiensis* | 117-1 | LC597586 | | HO | Lactic_min_117-1 |

| | | | | | | | |
|---|---|---|---|---|---|---|---|
| *Lacticaseibacillus* | *nasuensis* | | SU_18 | AB608051 | Unknown | HO | Lactic_nas_SU_18 |
| *Lacticaseibacillus* | *pantheris* | | JCM_12539 | NR_025189.1 | Unknown | HO | Lactic_pan_JCM_12539 |
| *Lacticaseibacillus* | *paracasei* | *paracasei* | JCM_8130 | NR_025880.1 | Nomadic | HO | Lactic_par_par_JCM_8130 |
| *Lacticaseibacillus* | *paracasei* | *tolerans* | JCM_1171 | AB181950 | Nomadic | HO | Lactic_par_tol_JCM_1171 |
| *Lacticaseibacillus* | *porcinae* | | R-42633 | HE616585 | Unknown | HO | Lactic_por_R-42633 |
| *Lacticaseibacillus* | *rhamnosus* | | ATCC_7469 | NR_043408.1 | Nomadic | HO | Lactic_rha_ATCC_7469 |
| *Lacticaseibacillus* | *saniviri* | | YIT_12363 | AB602569 | Unknown | HO | Lactic_san_YIT_12363 |
| *Lacticaseibacillus* | *sharpeae* | | JCM_1186 | NR_044711.2 | Unknown | HO | Lactic_sha_JCM_1186 |
| *Lacticaseibacillus* | *songhuajiangensis* | | 7-19 | HF679038 | Unknown | HO | Lactic_son_7-19 |
| *Lacticaseibacillus* | *suibinensis* | | 247-3 | MK110834 | Unknown | HO | Lactic_sui_247-3 |
| *Lacticaseibacillus* | *suilingensis* | | ZW152 | LC597590 | | HO | Lactic_sui_ZW152 |
| *Lacticaseibacillus* | *thailandensis* | | MCH5-2 | AB257863 | Unknown | HO | Lactic_tha_MCH5-2 |
| *Lacticaseibacillus* | *yichunensis* | | 33-1 | MK110845 | Unknown | HO | Lactic_yic_33-1 |
| *Lacticaseibacillus* | *zeae* | | ATCC_15820 | NR_037122.1 | | HO | Lactic_zeaeATCC_15820 |
| *Lacticaseibacillus* | *zhaodongensis* | | 1206-1 | LC508979 | | HO | Lactic_zha_1206-1 |
| *Lactiplantibacillus* | *argentoratensis* | | DK0_22 | AJ640078 | Nomadic | HO | Lactip_arg_DK0_22 |
| *Lactiplantibacillus* | *daoliensis* | | 116-1A | LC438516 | Nomadic | HO | Lactip_dao_116-1A |
| *Lactiplantibacillus* | *daowaiensis* | | 203-3 | LC438517 | Nomadic | HO | Lactip_dao_203-3 |
| *Lactiplantibacillus* | *dongliensis* | | 218-3 | LC438518 | Nomadic | HO | Lactip_don_218-3 |
| *Lactiplantibacillus* | *fabifermentans* | | DSM_21115 | AB626075 | Nomadic | HO | Lactip_fab_DSM_21115 |

| | | | | | | | |
|---|---|---|---|---|---|---|---|
| *Lactiplantibacillus* | *garii* | | FI11369 | MN817919 | | HO | Lactip_gar_FI11369 |
| *Lactiplantibacillus* | *herbarum* | | TCF032-E4 | KR706503 | Nomadic | HO | Lactip_her_TCF032-E4 |
| *Lactiplantibacillus* | *modestisalitolerans* | | NB446 | AB907192 | Nomadic | HO | Lactip_mod_NB446 |
| *Lactiplantibacillus* | *mudanjiangensis* | | 11050 | HF679037 | Nomadic | HO | Lactip_mud_11050 |
| *Lactiplantibacillus* | *nangangensis* | | 381-7 | LC438520 | Nomadic | HO | Lactip_nan_381-7 |
| *Lactiplantibacillus* | *paraplantarum* | | JCM_12533 | NR_025447.1 | Nomadic | HO | Lactip_par_JCM_12533 |
| *Lactiplantibacillus* | *pentosus* | | JCM_1558 | NR_029133.1 | Nomadic | HO | Lactip_pen_JCM_1558 |
| *Lactiplantibacillus* | *pingfangensis* | | 382-1 | LC438521 | Nomadic | HO | Lactip_pin_382-1 |
| *Lactiplantibacillus* | *plajomi* | | NB53 | AB907190 | Nomadic | HO | Lactip_pla_NB53 |
| *Lactiplantibacillus* | *plantarum* | *plantarum* | DSMZ_20174 | AJ965482 | Nomadic | HO | Lactip_pla_pla_DSMZ_20174 |
| *Lactiplantibacillus* | *songbeiensis* | | 398-2 | LC438523 | Nomadic | HO | Lactip_son_398-2 |
| *Lactiplantibacillus* | *xiangfangensis* | | 3.1.1 | HM443954 | Nomadic | HO | Lactip_xia_3.1.1 |
| *Lactobacillus* | *acetotolerans* | | ATCC_43578 | M58801 | Vertebrate | HO | Lactob_ace_ATCC_43578 |
| *Lactobacillus* | *acidophilus* | | ATCC_4356 | NR_043182.1 | Vertebrate | HO | Lactob_aci_ATCC_4356 |
| *Lactobacillus* | *amylolyticus* | | LA_5 | Y17361 | Vertebrate | HO | Lactob_amy_LA_5 |
| *Lactobacillus* | *amylovorus* | | ATCC_33620 | NR_043287.1 | Vertebrate | HO | Lactob_amy_ATCC_33620 |
| *Lactobacillus* | *apis* | | CCM_8403 | MT760199 | Insect | HO | Lactob_apisCCM_8403 |
| *Lactobacillus* | *bombicola* | | LMG_28288 | LK054485 | Insect | HO | Lactob_bom_LMG_28288 |
| *Lactobacillus* | *colini* | | 111144-L1 | KU161105 | Vertebrate | HO | Lactob_col_111144-L1 |

| | | | | | | | |
|---|---|---|---|---|---|---|---|
| *Lactobacillus* | *crispatus* | | strain_ATCC_33820 | NR_041800.1 | Vertebrate | HO | Lactob_cri_strain_ATCC_33820 |
| *Lactobacillus* | *delbrueckii* | *bulgaricus* | DSM20081(T) | NR_075019.1 | Vertebrate | HO | Lactob_del_bul_DSM20081(T) |
| *Lactobacillus* | *delbrueckii* | *delbrueckii* | DSMZ_20074 | NR_029106.1 | Vertebrate | HO | Lactob_del_del_DSMZ_20074 |
| *Lactobacillus* | *delbrueckii* | *indicus* | NCC_725 | AY421720 | Vertebrate | HO | Lactob_del_ind_NCC_725 |
| *Lactobacillus* | *delbrueckii* | *jakobsenii* | ZN7a-9 | JQ801728 | Vertebrate | HO | Lactob_del_jak_ZN7a-9 |
| *Lactobacillus* | *delbrueckii* | *lactis* | ATCC_12315 | NR_042728.1 | Vertebrate | HO | Lactob_del_lac_ATCC_12315 |
| *Lactobacillus* | *delbrueckii* | *sunkii* | YIT_11221 | AB641833 | Vertebrate | HO | Lactob_del_sun_YIT_11221 |
| *Lactobacillus* | *equicursoris* | | DI70 | AB290830 | Vertebrate | HO | Lactob_equ_DI70 |
| *Lactobacillus* | *fornicalis* | | JCM_12512 | NR_026509.1 | Vertebrate | HO | Lactob_for_JCM_12512 |
| *Lactobacillus* | *gallinarum* | | DSM_10532 | NR_042111.1 | Vertebrate | HO | Lactob_gal_DSM_10532 |
| *Lactobacillus* | *gasseri* | | JCM_1131 | NR_075051.2 | Vertebrate | HO | Lactob_gas_JCM_1131 |
| *Lactobacillus* | *gigeriorum* | | CRBIP_24.85 | NR_117057.1 | Vertebrate | HO | Lactob_gig_CRBIP_24.85 |
| *Lactobacillus* | *hamsteri* | | JCM_6256 | NR_025448.1 | Vertebrate | HO | Lactob_ham_JCM_6256 |
| *Lactobacillus* | *helsingborgensis* | | Bma5N | JX099553 | Insect | HO | Lactob_hel_Bma5N |
| *Lactobacillus* | *helveticus* | | ATCC_15009 | NR_042439.1 | Vertebrate | HO | Lactob_hel_ATCC_15009 |
| *Lactobacillus* | *hominis* | | CRBIP_24.179 | FR681902 | Vertebrate | HO | Lactob_hom_CRBIP_24.179 |
| *Lactobacillus* | *iners* | | CCUG_28746 | Y16329 | Vertebrate | HO | Lactob_ine_CCUG_2 |

| | | | | | | | |
|---|---|---|---|---|---|---|---|
| | | | | | | | 8746 |
| *Lactobacillus* | *intestinalis* | | JCM_7548 | NR_025449.1 | Vertebrate | HO | Lactob_int_JCM_7548 |
| *Lactobacillus* | *jensenii* | | ATCC_25258 | NR_025087.1 | Vertebrate | HO | Lactob_jen_ATCC_25258 |
| *Lactobacillus* | *johnsonii* | | DSM_10533 | NR_025273.1 | Vertebrate | HO | Lactob_joh_DSM_10533 |
| *Lactobacillus* | *kalixensis* | | Kx127A2 | AY253657 | Vertebrate | HO | Lactob_kal_Kx127A2 |
| *Lactobacillus* | *kefiranofaciens* | *kefiranofaciens* | ATCC_43761 | NR_042440.1 | Vertebrate | HO | Lactob_kef_kef_ATCC_43761 |
| *Lactobacillus* | *kefiranofaciens* | *kefirgranum* | ATCC_51647 | NR_042441.1 | Vertebrate | HO | Lactob_kef_kef_ATCC_51647 |
| *Lactobacillus* | *kimbladii* | | Hma2N | JX099549 | Insect | HO | Lactob_kim_Hma2N |
| *Lactobacillus* | *kitasatonis* | | JCM_1039 | NR_024813.1 | Vertebrate | HO | Lactob_kit_JCM_1039 |
| *Lactobacillus* | *kullabergensis* | | Biut2N | JX099550 | Insect | HO | Lactob_kul_Biut2N |
| *Lactobacillus* | *melliventris* | | Hma8N | JX099551 | Insect | HO | Lactob_mel_Hma8N |
| *Lactobacillus* | *mulieris* | | c10Ua161M | MK775269 | Vertebrate | HO | Lactob_mul_c10Ua161M |
| *Lactobacillus* | *panisapium* | | Bb_2-3 | KX447147 | Insect | HO | Lactob_pan_Bb_2-3 |
| *Lactobacillus* | *paragasseri* | | JCM_5343 | LC374363 | Vertebrate | HO | Lactob_par_JCM_5343 |
| *Lactobacillus* | *pasteurii* | | strain_1517 | FR681901 | Vertebrate | HO | Lactob_pas_strain_1517 |
| *Lactobacillus* | *porci* | | SG816 | MF346092 | Vertebrate | HO | Lactob_por_SG816 |
| *Lactobacillus* | *psittaci* | | JCM_11552 | AJ272391 | Vertebrate | HO | Lactob_psi_JCM_11552 |
| *Lactobacillus* | *rodentium* | | MYMRS/TLU1 | HQ851022 | Vertebrate | HO | Lactob_rod_MYMRS/TLU1 |
| *Lactobacillus* | *taiwanensis* | | BCRC_17755 | EU487512 | Vertebrate | HO | Lactob_tai_BCRC_1 |

| | | | | | | | |
|---|---|---|---|---|---|---|---|
| | | | | | | | 7755 |
| *Lactobacillus* | *ultunensis* | | Kx146C1 | AY253660 | Vertebrate | HO | Lactob_ult_Kx146C1 |
| *Lactobacillus* | *xujianguonis* | | HT111-2 | MK281589 | Vertebrate | HO | Lactob_xuj_HT111-2 |
| *Lapidilactobacillus* | *achengensis* | | 247-4 | MK110810 | Unknown | HO | Lapidi_ach_247-4 |
| *Lapidilactobacillus* | *bayanensis* | | 54-5 | MK110807 | Unknown | HO | Lapidi_bay_54-5 |
| *Lapidilactobacillus* | *concavus* | | JCM_14153 | LC258135 | Unknown | HO | Lapidi_con_JCM_14153 |
| *Lapidilactobacillus* | *dextrinicus* | | ATCC_33087 | NR_036861.1 | Unknown | HO | Lapidi_dex_ATCC_33087 |
| *Lapidilactobacillus* | *gannanensis* | | 143-1 | MK110813 | Unknown | HO | Lapidi_gan_143-1 |
| *Lapidilactobacillus* | *mulanensis* | | 143-6 | MK110808 | Unknown | HO | Lapidi_mul_143-6 |
| *Lapidilactobacillus* | *wuchangensis* | | 17-Apr | MK110811 | Unknown | HO | Lapidi_wuc_17-4 |
| *Latilactobacillus* | *curvatus* | | JCM_1096 | NR_042437.1 | Free Living | HO | Latila_cur_JCM_1096 |
| *Latilactobacillus* | *fuchuensis* | | JCM_11249 | NR_104976.1 | Free Living | HO | Latila_fuc_JCM_11249 |
| *Latilactobacillus* | *graminis* | | JCM_9503 | NR_042438.1 | Free Living | HO | Latila_gra_JCM_9503 |
| *Latilactobacillus* | *sakei* | *carnosus* | JCM_11031 | AY204892 | Free Living | HO | Latila_sak_car_JCM_11031 |
| *Latilactobacillus* | *sakei* | *sakei* | strain_ATCC_15521 | NR_042443.1 | Free Living | HO | Latila_sak_sak_strain_ATCC_15521 |
| *Lentilactobacillus* | *buchneri* | *silagei* | SG162 | LC507198 | | HE | Lentil_buc_sil_SG162 |
| *Lentilactobacillus* | *buchneri* | | JCM_1115 | NR_041293.1 | Free Living | HE | Lentil_buc_JCM_1115 |
| *Lentilactobacillus* | *curieae* | | S1L19 | JQ086550 | Free Living | HE | Lentil_cur_S1L19 |
| *Lentilactobacillus* | *diolivorans* | | JCM_12183 | NR_037004.1 | Free Living | HE | Lentil_dio_JCM_12183 |

| | | | | | | |
|---|---|---|---|---|---|---|
| *Lentilactobacillus* | *farraginis* | NRIC_0676 | AB262731 | Free Living | HE | Lentil_far_NRIC_0676 |
| *Lentilactobacillus* | *fungorum* | YK48G | LC383843 | | HE | Lentil_fun_YK48G |
| *Lentilactobacillus* | *hilgardii* | ATCC_8290 | NR_044708.2 | Free Living | HE | Lentil_hil_ATCC_8290 |
| *Lentilactobacillus* | *kefiri* | ATCC_35411 | NR_042230.1 | Free Living | HE | Lentil_kef_ATCC_35411 |
| *Lentilactobacillus* | *kisonensis* | YIT_11168 | AB366388 | Free Living | HE | Lentil_kis_YIT_11168 |
| *Lentilactobacillus* | *kosonis* | C06_No.73 | LC604908 | | HE | Lentil_kos_C06_No.73 |
| *Lentilactobacillus* | *kribbianus* | YH-lac9 | MN935214 | | HE | Lentil_kri_YH-lac9 |
| *Lentilactobacillus* | *otakiensis* | YIT_11163 | AB366386 | Free Living | HE | Lentil_ota_YIT_11163 |
| *Lentilactobacillus* | *parabuchneri* | JCM_12493 | NR_041294.1 | Free Living | HE | Lentil_par_JCM_12493 |
| *Lentilactobacillus* | *parafarraginis* | NRIC_0677 | AB262734 | Free Living | HE | Lentil_par_NRIC_0677 |
| *Lentilactobacillus* | *parakefiri* | JCM_8573 | NR_029039.1 | Free Living | HE | Lentil_par_JCM_8573 |
| *Lentilactobacillus* | *raoultii* | Marseille-P4006 | LT854294 | Free Living | HE | Lentil_rao_Marseille-P4006 |
| *Lentilactobacillus* | *rapi* | YIT_11204 | AB366389 | Free Living | HE | Lentil_rapi YIT_11204 |
| *Lentilactobacillus* | *senioris* | YIT_12364 | AB602570 | Unknown | HE | Lentil_sen_YIT_12364 |
| *Lentilactobacillus* | *sunkii* | YIT_11161 | AB366385 | Free Living | HE | Lentil_sun_YIT_11161 |
| *Leuconostoc* | *carnosum* | NCFB_2776 | NR_040811.1 | | HE | Leucon_car_NCFB_2776 |
| *Leuconostoc* | *citreum* | ATCC_49370 | AF111948 | | HE | Leucon_cit_ATCC_49370 |

| | | | | | | |
|---|---|---|---|---|---|---|
| *Leuconostoc* | *falkenbergense* | | LMG_10779 | HM443956 | HE | Leucon_fal_LMG_10779 |
| *Leuconostoc* | *fallax* | | DSM_20189 | NR_041830.1 | HE | Leucon_fal_DSM_20189 |
| *Leuconostoc* | *gasicomitatum* | | LMG_18811 | AF231131 | HE | Leucon_gas_LMG_18811 |
| *Leuconostoc* | *gelidum* | *aenigmaticum* | POUF4d | KF577569 | HE | Leucon_gel_aen_POUF4d |
| *Leuconostoc* | *gelidum* | | DSMZ_5578 | AF175402 | HE | Leucon_gel_DSMZ_5578 |
| *Leuconostoc* | *holzapfelii* | | LMG_23990 | AM600682 | HE | Leucon_hol_LMG_23990 |
| *Leuconostoc* | *inhae* | | IH003 | AF439560 | HE | Leucon_inh_IH003 |
| *Leuconostoc* | *kimchii* | | IH25 | AF173986 | HE | Leucon_kim_IH25 |
| *Leuconostoc* | *lactis* | | JCM_6123 | MT760652 | HE | Leucon_lac_JCM_6123 |
| *Leuconostoc* | *litchii* | | MB7 | LC259518 | HE | Leucon_lit_MB7 |
| *Leuconostoc* | *mesenteroides* | *cremoris* | NCFB_543 | AB023247 | HE | Leucon_mes_cre_NCFB_543 |
| *Leuconostoc* | *mesenteroides* | *dextranicum* | NCFB_529 | AB023244 | HE | Leucon_mes_dex_NCFB_529 |
| *Leuconostoc* | *mesenteroides* | *jonggajibkimchii* | DRC1506 | CP014611 | HE | Leucon_mes_jon_DRC1506 |
| *Leuconostoc* | *mesenteroides* | | ATCC_8293 | KX886793 | HE | Leucon_mes_ATCC_8293 |
| *Leuconostoc* | *miyukkimchii* | | M2 | HQ263024 | HE | Leucon_miy_M2 |
| *Leuconostoc* | *palmae* | | TMW_2.694 | AM940225 | HE | Leucon_pal_TMW_2.694 |
| *Leuconostoc* | *pseudomesenteroides* | | NCDO_768 | NR_040814.1 | HE | Leucon_pse_NCDO_768 |
| *Leuconostoc* | *rapi* | | LMG_27676 | HG515542 | HE | Leucon_rapiLMG_27 |

| | | | | | | |
|---|---|---|---|---|---|---|
| | | | | | | 676 |
| *Leuconostoc* | *suionicum* | LMG_8159 | HM443957 | | HE | Leucon_sui_LMG_8159 |
| *Levilactobacillus* | *acidifarinae* | CCM_7240 | NR_042242.1 | Free Living | HE | Levila_aci_CCM_7240 |
| *Levilactobacillus* | *andaensis* | 866-3 | LC597582 | | HE | Levila_and_866-3 |
| *Levilactobacillus* | *angrenensis* | M1530-1 | MK110858 | Free Living | HE | Levila_ang_M1530-1 |
| *Levilactobacillus* | *bambusae* | BCRC_80970 | KX400838 | Free Living | HE | Levila_bam_BCRC_80970 |
| *Levilactobacillus* | *brevis* | DSMZ_20054 | NR_044704.2 | Free Living | HE | Levila_bre_DSMZ_20054 |
| *Levilactobacillus* | *cerevisiae* | TUM_BP_140423000-2250 | KT445896 | Free Living | HE | Levila_cer_TUM_BP_140423000-2250 |
| *Levilactobacillus* | *enshiensis* | HBUAS57009 | MN082021 | Free Living | HE | Levila_ens_HBUAS57009 |
| *Levilactobacillus* | *fujinensis* | 218-6 | MK110865 | Free Living | HE | Levila_fuj_218-6 |
| *Levilactobacillus* | *fuyuanensis* | 244-4 | MK110862 | Free Living | HE | Levila_fuy_244-4 |
| *Levilactobacillus* | *hammesii* | TMW_1.1236 | AJ632219 | Free Living | HE | Levila_ham_TMW_1.1236 |
| *Levilactobacillus* | *huananensis* | 151-2B | MK110857 | Free Living | HE | Levila_hua_151-2B |
| *Levilactobacillus* | *koreensis* | DCY_50 | FJ904277 | Free Living | HE | Levila_kor_DCY_50 |
| *Levilactobacillus* | *lanxiensis* | 13B17 | LC597584 | | HE | Levila_lan_13B17 |
| *Levilactobacillus* | *lindianensis* | 220-4 | MK110856 | Free Living | HE | Levila_lin_220-4 |
| *Levilactobacillus* | *mulengensis* | 112-3 | MK110866 | Free Living | HE | Levila_mul_112-3 |
| *Levilactobacillus* | *namurensis* | LMG_23583 | AM259118 | Free Living | HE | Levila_nam_LMG_23583 |
| *Levilactobacillus* | *parabrevis* | LMG_11984 | AM158249 | Free Living | HE | Levila_par_LMG_11984 |

| | | | | | | |
|---|---|---|---|---|---|---|
| *Levilactobacillus* | *paucivorans* | TMW_1.1424 | FN185731 | Free Living | HE | Levila_pau_TMW_1.1424 |
| *Levilactobacillus* | *senmaizukei* | L13 | AB297927 | Free Living | HE | Levila_sen_L13 |
| *Levilactobacillus* | *spicheri* | LTH_5753 | AJ534844 | Free Living | HE | Levila_spi_LTH_5753 |
| *Levilactobacillus* | *suantsaii* | BCRC_12945 | MH730159 | Free Living | HE | Levila_sua_BCRC_12945 |
| *Levilactobacillus* | *suantsaiihabitans* | R19 | MH810313 | Free Living | HE | Levila_sua_R19 |
| *Levilactobacillus* | *tangyuanensis* | 137-3 | MK110859 | Free Living | HE | Levila_tan_137-3 |
| *Levilactobacillus* | *tongjiangensis* | 218-10 | MK110863 | Free Living | HE | Levila_ton_218-10 |
| *Levilactobacillus* | *wangkuiensis* | 6-5(1) | LC597583 | | HE | Levila_wan_6-5(1) |
| *Levilactobacillus* | *yonginensis* | THK-V8 | JN128640 | Free Living | HE | Levila_yon_THK-V8 |
| *Levilactobacillus* | *zymae* | CCM_7241 | MT760125 | Free Living | HE | Levila_zym_CCM_7241 |
| *Ligilactobacillus* | *acidipiscis* | JCM_10692 | NR_024718.1 | Unknown | HO | Ligila_aci_JCM_10692 |
| *Ligilactobacillus* | *agilis* | JCM_1187 | AB596945 | Unknown | HO | Ligila_agi_JCM_1187 |
| *Ligilactobacillus* | *animalis* | JCM_5670 | NR_041610.1 | Vertebrate | HO | Ligila_ani_JCM_5670 |
| *Ligilactobacillus* | *apodemi* | DSM_16634 | AB370874 | Vertebrate | HO | Ligila_apo_DSM_16634 |
| *Ligilactobacillus* | *araffinosus* | JCM_5667 | NR_104979.1 | Vertebrate | HO | Ligila_ara_JCM_5667 |
| *Ligilactobacillus* | *aviarius* | JCM_5666 | NR_044703.2 | Vertebrate | HO | Ligila_avi_JCM_5666 |
| *Ligilactobacillus* | *ceti* | strain_142-2 | AM292799 | Vertebrate | HO | Ligila_cetistrain_142-2 |
| *Ligilactobacillus* | *equi* | JCM_10991 | NR_028623.1 | Vertebrate | HO | Ligila_equiJCM_10991 |

| | | | | | | |
|---|---|---|---|---|---|---|
| *Ligilactobacillus* | *faecis* | AFL13-2 | NR_114391.1 | Vertebrate | HO | Ligila_fae_AFL13-2 |
| *Ligilactobacillus* | *hayakitensis* | KBL13 | AB267406 | Vertebrate | HO | Ligila_hay_KBL13 |
| *Ligilactobacillus* | *murinus* | JCM_1717 | NR_042231.1 | Vertebrate | HO | Ligila_mur_JCM_1717 |
| *Ligilactobacillus* | *pobuzihii* | NBRC_103219 | AB326358 | Unknown | HO | Ligila_pob_NBRC_103219 |
| *Ligilactobacillus* | *ruminis* | JCM_1152 | NR_041611.1 | Vertebrate | HO | Ligila_rum_JCM_1152 |
| *Ligilactobacillus* | *saerimneri* | GDA154 | AY255802 | Vertebrate | HO | Ligila_sae_GDA154 |
| *Ligilactobacillus* | *salitolerans* | YK43 | LC127508 | Unknown | HO | Ligila_sal_YK43 |
| *Ligilactobacillus* | *salivarius* | DSMZ_20555 | NR_028725.2 | Vertebrate | HO | Ligila_sal_DSMZ_20555 |
| *Limosilactobacillus* | *agrestis* | WF-MT5-A | MT823179 | | HE | Limosi_agr_WF-MT5-A |
| *Limosilactobacillus* | *albertensis* | Lr3000 | MT823181 | | HE | Limosi_alb_Lr3000 |
| *Limosilactobacillus* | *alvi* | R54 | HQ718585 | Vertebrate | HE | Limosi_alviR54 |
| *Limosilactobacillus* | *antri* | Kx146A4 | AY253659 | Vertebrate | HE | Limosi_ant_Kx146A4 |
| *Limosilactobacillus* | *balticus* | BG-AF3-A | MT823145 | | HE | Limosi_bal_BG-AF3-A |
| *Limosilactobacillus* | *caccae* | Marseille-P3519 | LT671596 | | HE | Limosi_cac_Marseille-P3519 |
| *Limosilactobacillus* | *caviae* | CCM_8609 | NR_157747.1 | Vertebrate | HE | Limosi_cav_CCM_8609 |
| *Limosilactobacillus* | *coleohominis* | JCM_11550 | NR_042436.1 | Vertebrate | HE | Limosi_col_JCM_11550 |
| *Limosilactobacillus* | *equigenerosi* | NRIC_0697 | AB288050 | Vertebrate | HE | Limosi_equ_NRIC_0697 |
| *Limosilactobacillus* | *fastidiosus* | WF-MO7-1 | MT823190 | | HE | Limosi_fas_WF-MO7-1 |

| | | | | | | | |
|---|---|---|---|---|---|---|---|
| *Limosilactobacillus* | *fermentum* | | strain_ATCC_14931 | NR_104927.1 | Unknown | HE | Limosi_fer_strain_ATCC_14931 |
| *Limosilactobacillus* | *frumenti* | | JCM_11122 | NR_025371.1 | Vertebrate | HE | Limosi_fru_JCM_11122 |
| *Limosilactobacillus* | *gastricus* | | strain_Kx156A7 | AY253658 | Vertebrate | HE | Limosi_gas_strain_Kx156A7 |
| *Limosilactobacillus* | *gorillae* | | KZ01 | AB904716 | Vertebrate | HE | Limosi_gor_KZ01 |
| *Limosilactobacillus* | *ingluviei* | | JCM_12531 | NR_028810.1 | Vertebrate | HE | Limosi_ing_JCM_12531 |
| *Limosilactobacillus* | *mucosae* | | JCM_12515 | NR_024994.1 | Vertebrate | HE | Limosi_muc_JCM_12515 |
| *Limosilactobacillus* | *oris* | | JCM_11028 | NR_026309.1 | Vertebrate | HE | Limosi_orisJCM_11028 |
| *Limosilactobacillus* | *panis* | | JCM_11053 | NR_026310.1 | Vertebrate | HE | Limosi_pan_JCM_11053 |
| *Limosilactobacillus* | *pontis* | | JCM_11051 | NR_036788.2 | Vertebrate | HE | Limosi_pon_JCM_11051 |
| *Limosilactobacillus* | *portuensis* | | c11Ua_112_M | MW016036 | | HE | Limosi_por_c11Ua_112_M |
| *Limosilactobacillus* | *reuteri* | *kinnaridis* | AP3 | MT823192 | | HE | Limosi_reu_kin_AP3 |
| *Limosilactobacillus* | *reuteri* | *murium* | lpuph1 | MW367781 | | HE | Limosi_reu_mur_lpuph1 |
| *Limosilactobacillus* | *reuteri* | *porcinus* | 3c6 | MW367860 | | HE | Limosi_reu_por_3c6 |
| *Limosilactobacillus* | *reuteri* | *rodentium* | 100-23 | MW367780 | | HE | Limosi_reu_rod_100-23 |
| *Limosilactobacillus* | *reuteri* | *suis* | ATCC_53608 | MW367845 | | HE | Limosi_reu_suisATCC_53608 |
| *Limosilactobacillus* | *reuteri* | | JCM_1112 | NR_025911.1 | Vertebrate | HE | Limosi_reu_JCM_1112 |
| *Limosilactobacillus* | *rudii* | | STM3_1 | MT823182 | | HE | Limosi_rud_STM3_ |

| | | | | | | |
|---|---|---|---|---|---|---|
| | | | | | | 1 |
| *Limosilactobacillus* | *secaliphilus* | TMW_1.1309 | AM279150 | Vertebrate | HE | Limosi_sec_TMW_1.1309 |
| *Limosilactobacillus* | *urinaemulieris* | c9Ua_26_M | MW016377 | | HE | Limosi_uri_c9Ua_26_M |
| *Limosilactobacillus* | *vaginalis* | JCM_9505 | NR_041796.1 | Vertebrate | HE | Limosi_vag_JCM_9505 |
| *Liquorilactobacillus* | *aquaticus* | strain_IMCC_1736 | DQ664203 | Free Living | HO | Liquor_aqu_strain_IMCC_1736 |
| *Liquorilactobacillus* | *cacaonum* | LMG_24285 | AM905389 | Free Living | HO | Liquor_cac_LMG_24285 |
| *Liquorilactobacillus* | *capillatus* | YIT_11306 | AB365976 | Free Living | HO | Liquor_cap_YIT_11306 |
| *Liquorilactobacillus* | *ghanensis* | L489 | DQ523489 | Free Living | HO | Liquor_gha_L489 |
| *Liquorilactobacillus* | *hordei* | UCC128 | EU074850 | Free Living | HO | Liquor_hor_UCC128 |
| *Liquorilactobacillus* | *mali* | JCM_1116 | NR_044709.2 | Free Living | HO | Liquor_maliJCM_1116 |
| *Liquorilactobacillus* | *nagelii* | JCM_12492 | NR_041007.1 | Free Living | HO | Liquor_nag_JCM_12492 |
| *Liquorilactobacillus* | *oeni* | strain_59b | AY681127 | Free Living | HO | Liquor_oenistrain_59b |
| *Liquorilactobacillus* | *satsumensis* | JCM_12392 | NR_028658.1 | Free Living | HO | Liquor_sat_JCM_12392 |
| *Liquorilactobacillus* | *sicerae* | CECT_8227 | HG794492 | Free Living | HO | Liquor_sic_CECT_8227 |
| *Liquorilactobacillus* | *sucicola* | NRIC_0736 | AB433982 | Free Living | HO | Liquor_suc_NRIC_0736 |
| *Liquorilactobacillus* | *uvarum* | strain_8 | NR_115308.1 | Free Living | HO | Liquor_uva_strain_8 |
| *Liquorilactobacillus* | *vini* | CECT_5924 | AJ576009 | Free Living | HO | Liquor_viniCECT_5924 |

| | | | | | | | |
|---|---|---|---|---|---|---|---|
| *Loigolactobacillus* | *backii* | | JCM_18665 | AB779648 | Unknown | HO | Loigol_bac_JCM_18665 |
| *Loigolactobacillus* | *bifermentans* | | JCM_1094 | NR_104926.1 | Unknown | HO | Loigol_bif_JCM_1094 |
| *Loigolactobacillus* | *binensis* | | 735-2 | LC438524 | Unknown | HO | Loigol_bin_735-2 |
| *Loigolactobacillus* | *coryniformis* | *coryniformis* | JCM_1164 | M58813 | Unknown | HO | Loigol_cor_cor_JCM_1164 |
| *Loigolactobacillus* | *coryniformis* | *torquens* | JCM_1166 | NR_029018.1 | Unknown | HO | Loigol_cor_tor_JCM_1166 |
| *Loigolactobacillus* | *iwatensis* | | strain_IWT246 | AB773428 | Unknown | HO | Loigol_iwa_strain_IWT246 |
| *Loigolactobacillus* | *jiayinensis* | | 257-1 | MK110846 | Unknown | HO | Loigol_jia_257-1 |
| *Loigolactobacillus* | *rennini* | | CECT_5922 | AJ576007 | Unknown | HO | Loigol_ren_CECT_5922 |
| *Loigolactobacillus* | *zhaoyuanensis* | | 187-3 | MK110851 | Unknown | HO | Loigol_zha_187-3 |
| Nicolia | spurrieriana | | SGEP1_A5 | OM986479 | Insect | HE | Nicol_spu_SGEP1_A5 |
| *Oenococcus* | *alcoholitolerans* | | UFRJ-M7.2.18 | HQ009794 | | HE | Oenoco_alc_UFRJ-M7.2.18 |
| *Oenococcus* | *kitaharae* | | NRIC_0645 | AB221475 | | HE | Oenoco_kit_NRIC_0645 |
| *Oenococcus* | *oeni* | | NCDO_1674 | NR_040810.1 | | HE | Oenoco_oeniNCDO_1674 |
| *Oenococcus* | *sicerae* | | UCMA_15228 | MH384882 | | HE | Oenoco_sic_UCMA_15228 |
| *Paralactobacillus* | *selangorensis* | | LMG_17710 | NR_024885.1 | Unknown | HO | Parala_sel_LMG_17710 |
| *Paucilactobacillus* | *hokkaidonensis* | | LOOC260 | AB721549 | Free Living | HE | Paucil_hok_LOOC260 |
| *Paucilactobacillus* | *kaifaensis* | | 778-3 | LC438525 | Free Living | HE | Paucil_kai_778-3 |
| *Paucilactobacillus* | *nenjiangensis* | | 11102 | HF679039 | Free Living | HE | Paucil_nen_11102 |

| | | | | | | |
|---|---|---|---|---|---|---|
| *Paucilactobacillus* | *oligofermentans* | AMKR18 | AY733084 | Free Living | HE | Paucil_oli_AMKR18 |
| *Paucilactobacillus* | *suebicus* | JCM_9504 | NR_042190.1 | Free Living | HE | Paucil_sue_JCM_9504 |
| *Paucilactobacillus* | *vaccinostercus* | DSM_20634 | NR_042233.1 | Free Living | HE | Paucil_vac_DSM_20634 |
| *Paucilactobacillus* | *wasatchensis* | WDC04 | AWTT01000084 | Free Living | HE | Paucil_was_WDC04 |
| *Pediococcus* | *acidilactici* | DSMZ_20284 | NR_041640.1 | | HO | Pedioc_aci_DSMZ_20284 |
| *Pediococcus* | *argentinicus* | LMG_23999 | AM709786 | | HO | Pedioc_arg_LMG_23999 |
| *Pediococcus* | *cellicola* | JCM_14152 | LC258134 | | HO | Pedioc_cel_JCM_14152 |
| *Pediococcus* | *claussenii* | DSM_14800 | AJ621555 | | HO | Pedioc_cla_DSM_14800 |
| *Pediococcus* | *damnosus* | ATCC_29358 | NR_042087.1 | | HO | Pedioc_dam_ATCC_29358 |
| *Pediococcus* | *ethanolidurans* | Z-9 | AY956789 | | HO | Pedioc_eth_Z-9 |
| *Pediococcus* | *inopinatus* | ATCC_49902 | NR_025388.1 | | HO | Pedioc_ino_ATCC_49902 |
| *Pediococcus* | *parvulus* | JCM_5889 | NR_029136.1 | | HO | Pedioc_par_JCM_5889 |
| *Pediococcus* | *pentosaceus* | ATCC_33314 | NR_042058.1 | | HO | Pedioc_pen_ATCC_33314 |
| *Pediococcus* | *siamensis* | MCH3-2 | AB258357 | | HO | Pedioc_sia_MCH3-2 |
| *Pediococcus* | *stilesii* | LMG_23082 | AJ973157 | | HO | Pedioc_sti_LMG_23082 |
| *Periweissella* | *beninensis* | 2L24P13 | EU439435 | | HO | Periwe_ben_2L24P13 |
| *Periweissella* | *cryptocerci* | 26KH-42 | MK395366 | | HO | Periwe_cry_26KH-42 |

| | | | | | | |
|---|---|---|---|---|---|---|
| *Periweissella* | *fabalis* | LMG_26217 | HE576795 | | HO | Periwe_fab_LMG_26217 |
| *Periweissella* | *fabaria* | 257 | FM179678 | | HO | Periwe_fab_257 |
| *Periweissella* | *ghanensis* | LMG_24286 | AM882997 | | HO | Periwe_gha_LMG_24286 |
| *Schleiferilactobacillus* | *harbinensis* | NBRC_100982 | AB681318 | Free Living | HO | Schlei_har_NBRC_100982 |
| *Schleiferilactobacillus* | *perolens* | JCM_12534 | NR_029360.1 | Free Living | HO | Schlei_per_JCM_12534 |
| *Schleiferilactobacillus* | *shenzhenensis* | KACC_16878 | JX523627 | Free Living | HO | Schlei_she_KACC_16878 |
| *Secundilactobacillus* | *collinoides* | JCM_1123 | NR_024645.1 | Free Living | HE | Secund_col_JCM_1123 |
| *Secundilactobacillus* | *folii* | CRM56-3 | LC511609 | | HE | Secund_fol_CRM56-3 |
| *Secundilactobacillus* | *hailunensis* | 887-11 | LC473139 | | HE | Secund_hai_887-11 |
| *Secundilactobacillus* | *kimchicus* | DCY_51 | EU678893 | Free Living | HE | Secund_kim_DCY_51 |
| *Secundilactobacillus* | *malefermentans* | JCM_12497 | NR_042442.1 | Free Living | HE | Secund_mal_JCM_12497 |
| *Secundilactobacillus* | *mixtipabuli* | IWT30 | AB894864 | Free Living | HE | Secund_mix_IWT30 |
| *Secundilactobacillus* | *odoratitofui* | YIT_11304 | AB365975 | Free Living | HE | Secund_odo_YIT_11304 |
| *Secundilactobacillus* | *oryzae* | SG293 | AB731660 | Free Living | HE | Secund_ory_SG293 |
| *Secundilactobacillus* | *paracollinoides* | JCM_11969 | NR_042322.1 | Free Living | HE | Secund_par_JCM_11969 |
| *Secundilactobacillus* | *pentosiphilus* | IWT25 | LC085284 | Free Living | HE | Secund_pen_IWT25 |
| *Secundilactobacillus* | *silagei* | strain_IWT126 | AB786910 | Free Living | HE | Secund_sil_strain_IWT126 |
| *Secundilactobacillus* | *silagincola* | IWT5 | LC085283 | Free Living | HE | Secund_sil_IWT5 |

| | | | | | | |
|---|---|---|---|---|---|---|
| *Secundilactobacillus* | *similis* | JCM_2765 | NR_112645.1 | Free Living | HE | Secund_sim_JCM_2765 |
| *Secundilactobacillus* | *yichangensis* | F79-211-2 | LC597581 | | HE | Secund_yic_F79-211-2 |
| *Weissella* | *bombi* | LMG_28290 | LK054487 | | HE | Weisse_bom_LMG_28290 |
| *Weissella* | *ceti* | strain_1119-1A-09 | FN813251 | | HE | Weisse_cetistrain_1119-1A-09 |
| *Weissella* | *cibaria* | LMG_17699 | AJ295989 | | HE | Weisse_cib_LMG_17699 |
| *Weissella* | *coleopterorum* | HDW19 | MN099422 | | HE | Weisse_col_HDW19 |
| *Weissella* | *confusa* | JCM_1093 | AB023241 | Unknown | HE | Weisse_con_JCM_1093 |
| *Weissella* | *diestrammenae* | ORY33 | JQ646523 | | HE | Weisse_die_ORY33 |
| *Weissella* | *halotolerans* | NRIC_1627 | AB022926 | Unknown | HE | Weisse_hal_NRIC_1627 |
| *Weissella* | *hellenica* | NCFB_2973 | X95981 | | HE | Weisse_hel_NCFB_2973 |
| *Weissella* | *kandleri* | NRIC_1628 | AB022922 | Unknown | HE | Weisse_kan_NRIC_1628 |
| *Weissella* | *koreensis* | JCM_11263 | LC145573 | | HE | Weisse_kor_JCM_11263 |
| *Weissella* | *minor* | NRIC_1625 | AB022920 | Unknown | HE | Weisse_min_NRIC_1625 |
| *Weissella* | *muntiaci* | 8_H-2 | MK774696 | | HE | Weisse_mun_8_H-2 |
| *Weissella* | *oryzae* | SG25 | AB690345 | | HE | Weisse_ory_SG25 |
| *Weissella* | *paramesenteroides* | NCDO_803 | NR_104568.1 | | HE | Weisse_par_NCDO_803 |
| *Weissella* | *sagaensis* | X0750 | LC438526 | | HE | Weisse_sag_X0750 |
| *Weissella* | *salipiscis* | FS55-1 | AB257595 | | HE | Weisse_sal_FS55-1 |

| | | | | | | |
|---|---|---|---|---|---|---|
| *Weissella* | *soli* | JCM_12536 | LC258131 | | HE | Weisse_soliJCM_12536 |
| *Weissella* | *thailandensis* | PCU_210 | NR_040822.1 | | HE | Weisse_tha_PCU_210 |
| *Weissella* | *uvarum* | B18NM42 | KF999666 | | HE | Weisse_uva_B18NM42 |
| *Weissella* | *viridescens* | NCDO_1655 | X52568 | Unknown | HE | Weisse_vir_NCDO_1655 |
| *Carnobacterium* | *maltaromaticum* | ATCC_27865 | NR_044710.2 | Unknown | HO | Carnob_ma |

**Supplementary Figure 1**. Maximum-likelihood phylogenetic tree showing the evolutionary relationships among sequences of the V1-V3 region of the 16S rRNA gene for type strains of species belonging to the family *Lactobacillaceae.* Leaves of species belonging to the same genus share the same color. The scale (number of nucleotide substitutions per nucleotide) is shown.

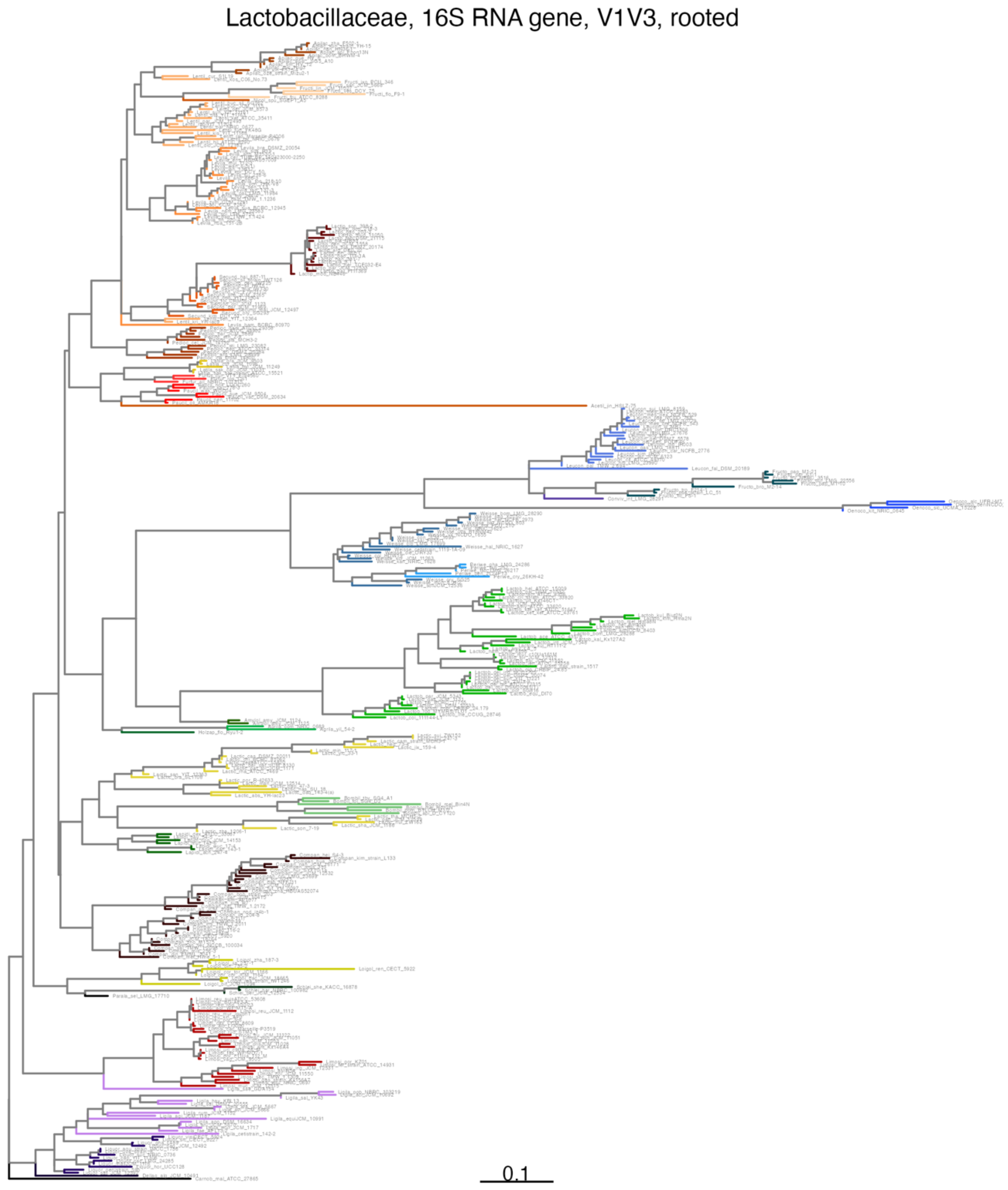

**Supplementary Figure 2**. Maximum-likelihood phylogenetic tree showing the evolutionary relationships among sequences of the V3-V4 region of the 16S rRNA gene for type strains of species belonging to the family *Lactobacillaceae.* Leaves of species belonging to the same genus share the same color. The scale (number of nucleotide substitutions per nucleotide) is shown.

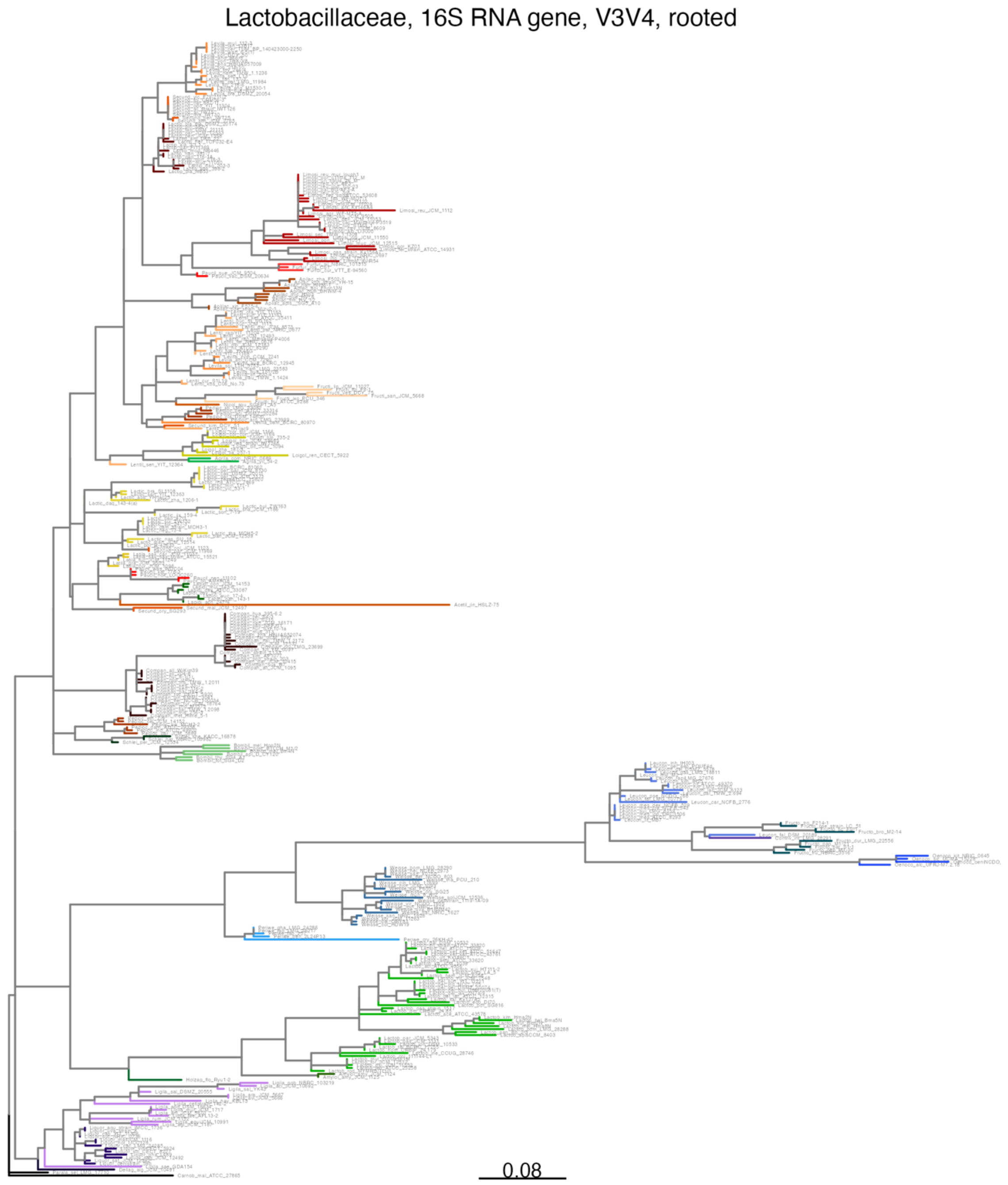

**Supplementary Figure 3**. Maximum-likelihood phylogenetic tree showing the evolutionary relationships among sequences of the V4 region of the 16S rRNA gene for type strains of species belonging to the family *Lactobacillaceae.* Leaves of species belonging to the same genus share the same color. The scale (number of nucleotide substitutions per nucleotide) is shown.

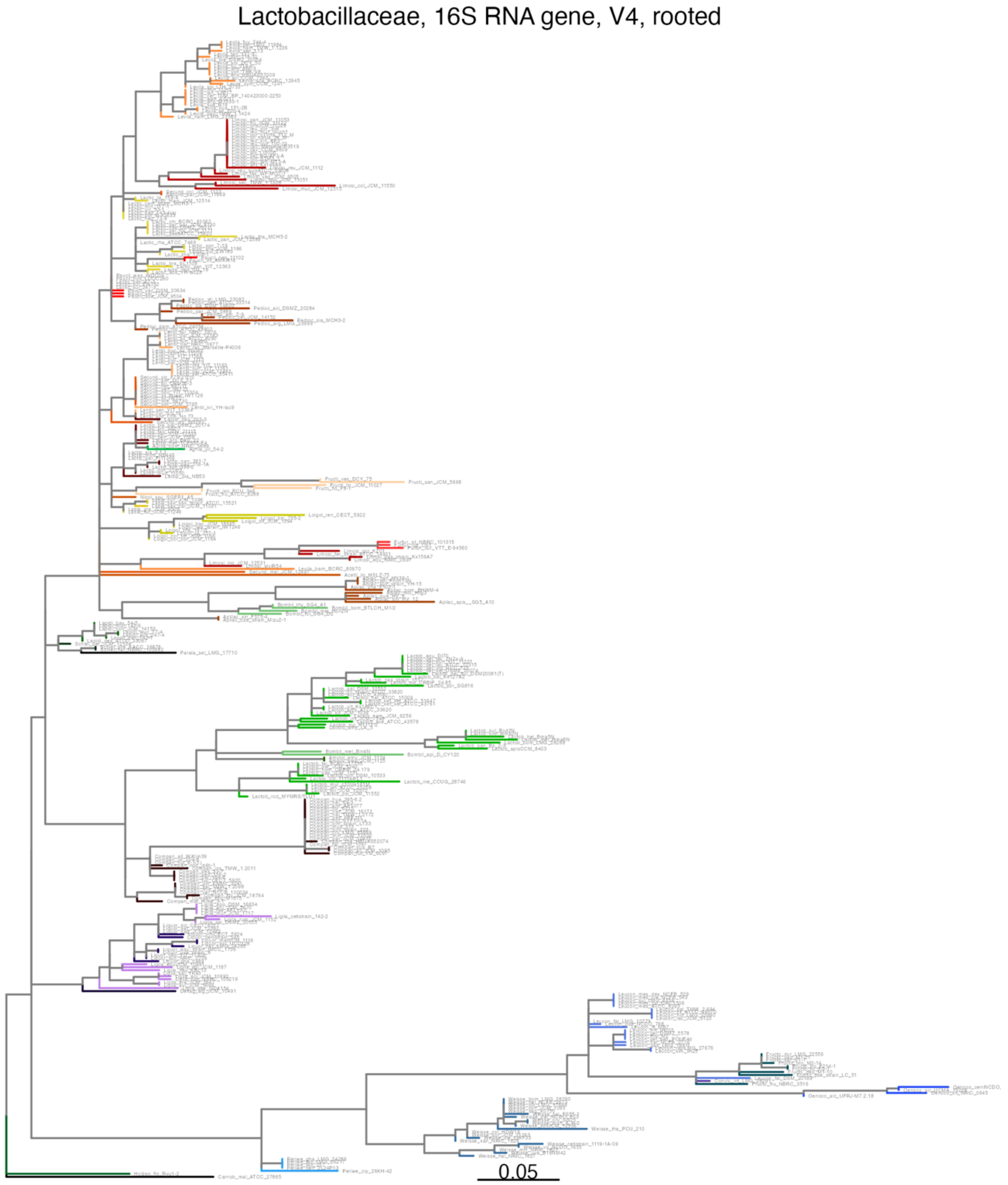

**Supplementary Table 2.** Sensitivity, specificity, positive predictive value and negative predictive value for confusion matrices for taxonomic assignment of 16S rRNA gene sequences (full sequence and V1-V3, V3-V4, V4 regions) of the type strains for genera of family *Lactobacillaceae*.

| region | *class* | sensitivity | specificity | pos_pred_value | neg_pred_value |
|---|---|---|---|---|---|
| FL | *Acetilactobacillus* | 0.000 | 1.000 | NA | 0.997 |
| FL | *Agrilactobacillus* | 1.000 | 1.000 | 1.000 | 1.000 |
| FL | *Amylolactobacillus* | 1.000 | 1.000 | 1.000 | 1.000 |
| FL | *Apilactobacillus* | 1.000 | 0.997 | 0.917 | 1.000 |
| FL | *Bombilactobacillus* | 1.000 | 1.000 | 1.000 | 1.000 |
| FL | *Companilactobacillus* | 1.000 | 1.000 | 1.000 | 1.000 |
| FL | *Convivina* | 0.000 | 1.000 | NA | 0.997 |
| FL | *Dellaglioa* | 1.000 | 1.000 | 1.000 | 1.000 |
| FL | *Fructilactobacillus* | 1.000 | 1.000 | 1.000 | 1.000 |
| FL | *Fructobacillus* | 1.000 | 1.000 | 1.000 | 1.000 |
| FL | *Furfurilactobacillus* | 1.000 | 1.000 | 1.000 | 1.000 |
| FL | *Holzapfelia* | 1.000 | 1.000 | 1.000 | 1.000 |
| FL | *Lacticaseibacillus* | 1.000 | 1.000 | 1.000 | 1.000 |
| FL | *Lactiplantibacillus* | 1.000 | 1.000 | 1.000 | 1.000 |
| FL | *Lactobacillus* | 1.000 | 0.997 | 0.978 | 1.000 |
| FL | *Lapidilactobacillus* | 1.000 | 1.000 | 1.000 | 1.000 |
| FL | *Latilactobacillus* | 1.000 | 1.000 | 1.000 | 1.000 |
| FL | *Lentilactobacillus* | 1.000 | 1.000 | 1.000 | 1.000 |
| FL | *Leuconostoc* | 1.000 | 0.997 | 0.955 | 1.000 |
| FL | *Levilactobacillus* | 0.963 | 1.000 | 1.000 | 0.997 |
| FL | *Ligilactobacillus* | 1.000 | 1.000 | 1.000 | 1.000 |
| FL | *Limosilactobacillus* | 1.000 | 1.000 | 1.000 | 1.000 |
| FL | *Liquorilactobacillus* | 1.000 | 1.000 | 1.000 | 1.000 |
| FL | *Loigolactobacillus* | 1.000 | 1.000 | 1.000 | 1.000 |
| FL | *Nicolia* | 0.000 | 1.000 | NA | 0.997 |
| FL | *Oenococcus* | 1.000 | 1.000 | 1.000 | 1.000 |
| FL | *Paralactobacillus* | 1.000 | 1.000 | 1.000 | 1.000 |
| FL | *Paucilactobacillus* | 1.000 | 1.000 | 1.000 | 1.000 |
| FL | *Pediococcus* | 1.000 | 1.000 | 1.000 | 1.000 |
| FL | *Periweissella* | 0.000 | 1.000 | NA | 0.987 |
| FL | *Schleiferilactobacillus* | 1.000 | 1.000 | 1.000 | 1.000 |
| FL | *Secundilactobacillus* | 1.000 | 0.997 | 0.933 | 1.000 |
| FL | *Weissella* | 1.000 | 0.986 | 0.800 | 1.000 |
| V1V3 | *Acetilactobacillus* | 0.000 | 1.000 | NA | 0.997 |
| V1V3 | *Agrilactobacillus* | 1.000 | 1.000 | 1.000 | 1.000 |
| V1V3 | *Amylolactobacillus* | 1.000 | 1.000 | 1.000 | 1.000 |
| V1V3 | *Apilactobacillus* | 1.000 | 0.997 | 0.917 | 1.000 |
| V1V3 | *Bombilactobacillus* | 1.000 | 1.000 | 1.000 | 1.000 |

| | | | | | |
|---|---|---|---|---|---|
| V1V3 | *Companilactobacillus* | 1.000 | 1.000 | 1.000 | 1.000 |
| V1V3 | *Convivina* | 0.000 | 1.000 | NA | 0.997 |
| V1V3 | *Dellaglioa* | 1.000 | 1.000 | 1.000 | 1.000 |
| V1V3 | *Fructilactobacillus* | 1.000 | 1.000 | 1.000 | 1.000 |
| V1V3 | *Fructobacillus* | 1.000 | 1.000 | 1.000 | 1.000 |
| V1V3 | *Furfurilactobacillus* | 1.000 | 1.000 | 1.000 | 1.000 |
| V1V3 | *Holzapfelia* | 1.000 | 1.000 | 1.000 | 1.000 |
| V1V3 | *HT002* | NA | 0.981 | NA | NA |
| V1V3 | *Lacticaseibacillus* | 1.000 | 1.000 | 1.000 | 1.000 |
| V1V3 | *Lactiplantibacillus* | 1.000 | 1.000 | 1.000 | 1.000 |
| V1V3 | *Lactobacillus* | 1.000 | 0.997 | 0.978 | 1.000 |
| V1V3 | *Lapidilactobacillus* | 1.000 | 1.000 | 1.000 | 1.000 |
| V1V3 | *Latilactobacillus* | 1.000 | 1.000 | 1.000 | 1.000 |
| V1V3 | *Lentilactobacillus* | 1.000 | 1.000 | 1.000 | 1.000 |
| V1V3 | *Leuconostoc* | 1.000 | 0.997 | 0.955 | 1.000 |
| V1V3 | *Levilactobacillus* | 0.963 | 1.000 | 1.000 | 0.997 |
| V1V3 | *Ligilactobacillus* | 1.000 | 1.000 | 1.000 | 1.000 |
| V1V3 | *Limosilactobacillus* | 0.767 | 1.000 | 1.000 | 0.980 |
| V1V3 | *Liquorilactobacillus* | 1.000 | 1.000 | 1.000 | 1.000 |
| V1V3 | *Loigolactobacillus* | 1.000 | 1.000 | 1.000 | 1.000 |
| V1V3 | *Nicolia* | 0.000 | 1.000 | NA | 0.997 |
| V1V3 | *Oenococcus* | 1.000 | 1.000 | 1.000 | 1.000 |
| V1V3 | *Paralactobacillus* | 1.000 | 1.000 | 1.000 | 1.000 |
| V1V3 | *Paucilactobacillus* | 1.000 | 1.000 | 1.000 | 1.000 |
| V1V3 | *Pediococcus* | 1.000 | 1.000 | 1.000 | 1.000 |
| V1V3 | *Periweissella* | 0.000 | 1.000 | NA | 0.987 |
| V1V3 | *Schleiferilactobacillus* | 1.000 | 1.000 | 1.000 | 1.000 |
| V1V3 | *Secundilactobacillus* | 1.000 | 0.997 | 0.933 | 1.000 |
| V1V3 | *Weissella* | 1.000 | 0.986 | 0.800 | 1.000 |
| V3V4 | *Acetilactobacillus* | NA | 1.000 | NA | NA |
| V3V4 | *Agrilactobacillus* | 1.000 | 1.000 | 1.000 | 1.000 |
| V3V4 | *Amylolactobacillus* | 1.000 | 1.000 | 1.000 | 1.000 |
| V3V4 | *Apilactobacillus* | 1.000 | 0.997 | 0.917 | 1.000 |
| V3V4 | *Bombilactobacillus* | 1.000 | 1.000 | 1.000 | 1.000 |
| V3V4 | *Companilactobacillus* | 1.000 | 1.000 | 1.000 | 1.000 |
| V3V4 | *Convivina* | 0.000 | 1.000 | NA | 0.997 |
| V3V4 | *Dellaglioa* | 1.000 | 1.000 | 1.000 | 1.000 |
| V3V4 | *Fructilactobacillus* | 1.000 | 1.000 | 1.000 | 1.000 |
| V3V4 | *Fructobacillus* | 1.000 | 1.000 | 1.000 | 1.000 |
| V3V4 | *Furfurilactobacillus* | 1.000 | 1.000 | 1.000 | 1.000 |
| V3V4 | *Holzapfelia* | 1.000 | 1.000 | 1.000 | 1.000 |
| V3V4 | *Lacticaseibacillus* | 0.880 | 1.000 | 1.000 | 0.991 |
| V3V4 | *Lactiplantibacillus* | 1.000 | 1.000 | 1.000 | 1.000 |

| | | | | | |
|---|---|---|---|---|---|
| V3V4 | *Lactobacillus* | 1.000 | 1.000 | 1.000 | 1.000 |
| V3V4 | *Lapidilactobacillus* | 1.000 | 1.000 | 1.000 | 1.000 |
| V3V4 | *Latilactobacillus* | 1.000 | 1.000 | 1.000 | 1.000 |
| V3V4 | *Lentilactobacillus* | 0.947 | 1.000 | 1.000 | 0.997 |
| V3V4 | *Leuconostoc* | 1.000 | 0.997 | 0.955 | 1.000 |
| V3V4 | *Levilactobacillus* | 0.963 | 1.000 | 1.000 | 0.997 |
| V3V4 | *Ligilactobacillus* | 1.000 | 1.000 | 1.000 | 1.000 |
| V3V4 | *Limosilactobacillus* | 0.679 | 1.000 | 1.000 | 0.975 |
| V3V4 | *Liquorilactobacillus* | 1.000 | 1.000 | 1.000 | 1.000 |
| V3V4 | *Loigolactobacillus* | 1.000 | 1.000 | 1.000 | 1.000 |
| V3V4 | *Nicolia* | 0.000 | 1.000 | NA | 0.997 |
| V3V4 | *Oenococcus* | 1.000 | 1.000 | 1.000 | 1.000 |
| V3V4 | *Paralactobacillus* | 1.000 | 1.000 | 1.000 | 1.000 |
| V3V4 | *Paucilactobacillus* | 1.000 | 1.000 | 1.000 | 1.000 |
| V3V4 | *Pediococcus* | 1.000 | 1.000 | 1.000 | 1.000 |
| V3V4 | *Periweissella* | 0.000 | 1.000 | NA | 0.987 |
| V3V4 | *Schleiferilactobacillus* | 1.000 | 1.000 | 1.000 | 1.000 |
| V3V4 | *Secundilactobacillus* | 1.000 | 0.986 | 0.737 | 1.000 |
| V3V4 | *Weissella* | 1.000 | 0.986 | 0.800 | 1.000 |
| V4 | *Acetilactobacillus* | NA | 1.000 | NA | NA |
| V4 | *Agrilactobacillus* | 0.000 | 1.000 | NA | 0.997 |
| V4 | *Amylolactobacillus* | 1.000 | 1.000 | 1.000 | 1.000 |
| V4 | *Apilactobacillus* | 1.000 | 1.000 | 1.000 | 1.000 |
| V4 | *Bombilactobacillus* | 1.000 | 1.000 | 1.000 | 1.000 |
| V4 | *Companilactobacillus* | 1.000 | 1.000 | 1.000 | 1.000 |
| V4 | *Convivina* | 0.000 | 1.000 | NA | 0.997 |
| V4 | *Dellaglioa* | 1.000 | 1.000 | 1.000 | 1.000 |
| V4 | *Fructilactobacillus* | 1.000 | 1.000 | 1.000 | 1.000 |
| V4 | *Fructobacillus* | 1.000 | 0.994 | 0.818 | 1.000 |
| V4 | *Furfurilactobacillus* | 1.000 | 1.000 | 1.000 | 1.000 |
| V4 | *Holzapfelia* | 1.000 | 1.000 | 1.000 | 1.000 |
| V4 | HT002 | NA | 0.956 | NA | NA |
| V4 | *Lacticaseibacillus* | 0.850 | 1.000 | 1.000 | 0.991 |
| V4 | *Lactiplantibacillus* | 1.000 | 0.997 | 0.941 | 1.000 |
| V4 | *Lactobacillus* | 1.000 | 1.000 | 1.000 | 1.000 |
| V4 | *Lapidilactobacillus* | 1.000 | 1.000 | 1.000 | 1.000 |
| V4 | *Latilactobacillus* | 1.000 | 1.000 | 1.000 | 1.000 |
| V4 | *Lentilactobacillus* | 1.000 | 1.000 | 1.000 | 1.000 |
| V4 | *Leuconostoc* | 0.952 | 1.000 | 1.000 | 0.997 |
| V4 | *Levilactobacillus* | 0.963 | 1.000 | 1.000 | 0.997 |
| V4 | *Ligilactobacillus* | 1.000 | 1.000 | 1.000 | 1.000 |
| V4 | *Limosilactobacillus* | 0.407 | 1.000 | 1.000 | 0.955 |
| V4 | *Liquorilactobacillus* | 1.000 | 1.000 | 1.000 | 1.000 |

| | | | | | |
|---|---|---|---|---|---|
| V4 | *Loigolactobacillus* | 1.000 | 1.000 | 1.000 | 1.000 |
| V4 | *Nicolia* | NA | 1.000 | NA | NA |
| V4 | *Oenococcus* | 1.000 | 1.000 | 1.000 | 1.000 |
| V4 | *Paralactobacillus* | 1.000 | 1.000 | 1.000 | 1.000 |
| V4 | *Paucilactobacillus* | 1.000 | 0.992 | 0.700 | 1.000 |
| V4 | *Pediococcus* | 1.000 | 1.000 | 1.000 | 1.000 |
| V4 | *Periweissella* | 0.000 | 1.000 | NA | 0.989 |
| V4 | *Schleiferilactobacillus* | 1.000 | 1.000 | 1.000 | 1.000 |
| V4 | *Secundilactobacillus* | 1.000 | 0.997 | 0.933 | 1.000 |
| V4 | *Weissella* | 1.000 | 0.988 | 0.833 | 1.000 |

**Supplementary Figure 4.** Distribution of the within-genus proportion of matching species identification for full-length sequences of the 16S rRNA gene and V1-V3, V3-V4, V4 for type strains of the species belonging to family *Lactobacillaceae*.

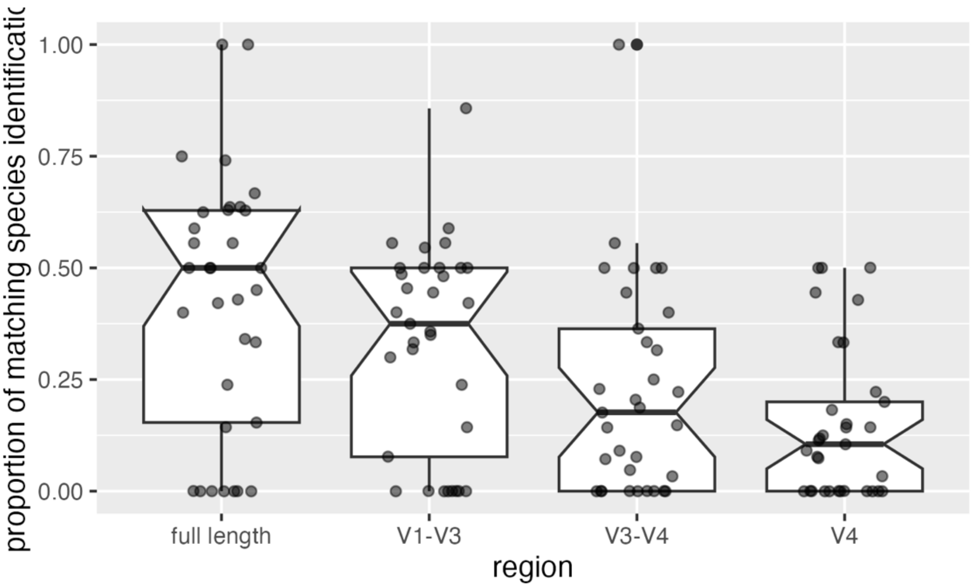

**Supplementary Table 3**. List of L1 categories in the EFSA FoodEx2 classification.

| L1 |
|---|
| Grains and grain-based products |
| Vegetables and vegetable products |
| Starchy roots or tubers and products thereof, sugar plants |
| Legumes, nuts, oilseeds and spices |
| Fruit and fruit products |
| Meat and meat products |
| Fish, seafood, amphibians, reptiles and invertebrates |
| Milk and dairy products |
| Eggs and egg products |
| Sugar and similar, confectionery and water-based sweet desserts |
| Animal and vegetable fats and oils and primary derivatives thereof |
| Fruit and vegetable juices and nectars (including concentrates) |
| Water and water-based beverages |
| Alcoholic beverages |
| Coffee, cocoa, tea and infusions |
| Food products for young population |
| Products for non-standard diets, food imitates and food supplements |
| Composite dishes |
| Seasoning, sauces and condiments |
| Major isolated ingredients, additives, flavours, baking and processing aids |
| Other ingredients |

**Supplementary Figure 5**. Box and jitter showing the distribution of relative abundance of *Lacticaseibacillus* sequences (only sequences for the V1-V3 and V3-V4 regions with length >350 bp were selected) in food samples from FoodMicrobionet. The individual points show the abundance in samples and their colour shows the nature of samples (unspoiled, fermented, spoiled or both).

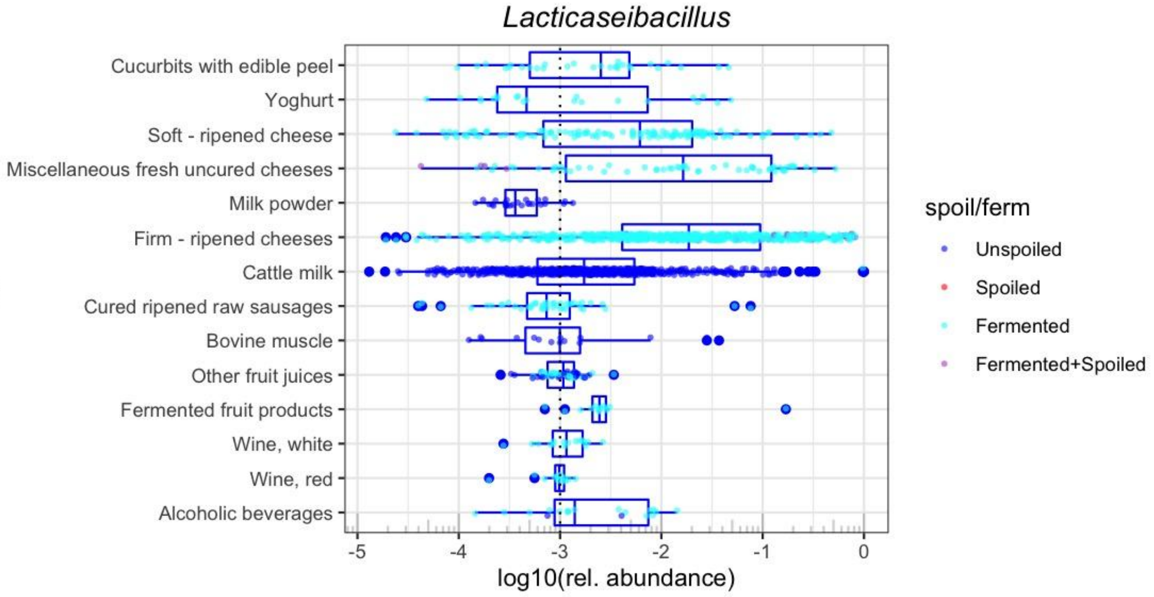


**Supplementary Figure 6**. Box and jitter showing the distribution of relative abundance of *Holzapfelia* sequences (only sequences for the V1-V3 and V3-V4 regions with length >350 bp were selected) in food samples from FoodMicrobionet. The individual points show the abundance in samples and their colour shows the nature of samples (unspoiled, fermented, spoiled or both).

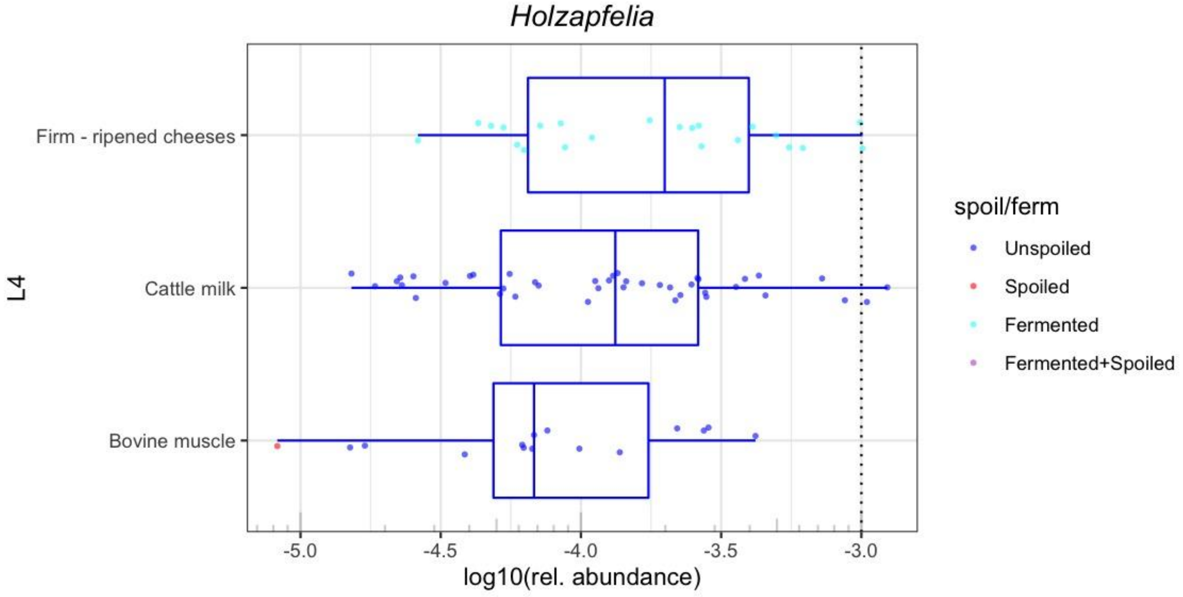

**Supplementary Figure 7**. Box and jitter showing the distribution of relative abundance of *Schleferilactobacillus* sequences (only sequences for the V1-V3 and V3-V4 regions with length >350 bp were selected) in food samples from FoodMicrobionet. The individual points show the abundance in samples and their colour shows the nature of samples (unspoiled, fermented, spoiled or both).

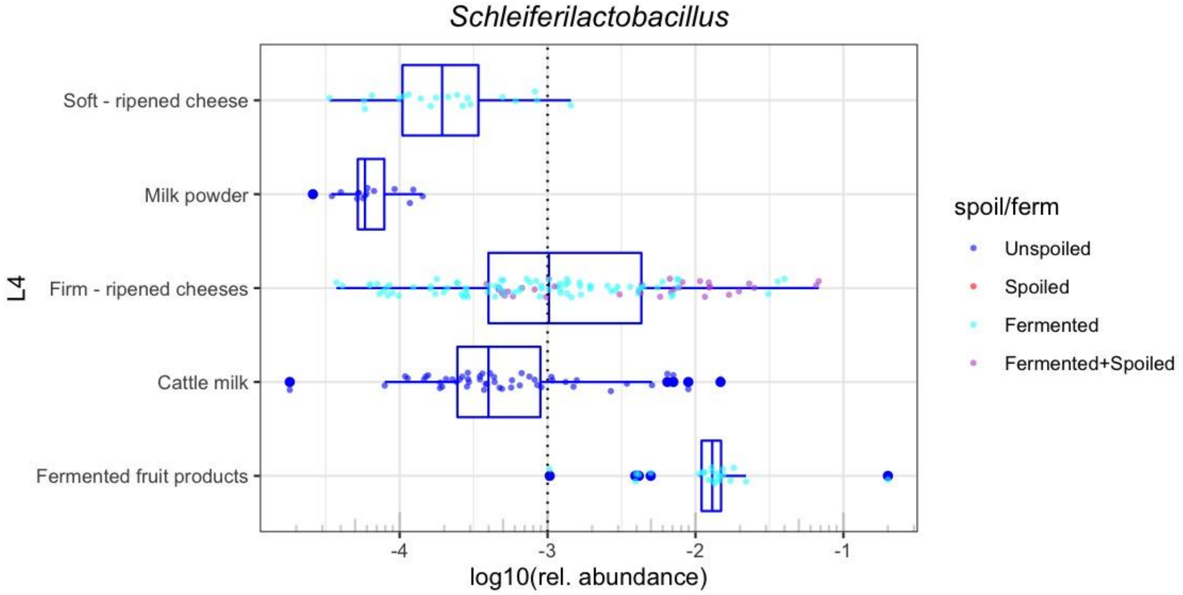


**Supplementary Figure 8**. Box and jitter showing the distribution of relative abundance of *Liquorilactobacillus* sequences (only sequences for the V1-V3 and V3-V4 regions with length >350 bp were selected) in food samples from FoodMicrobionet. The individual points show the abundance in samples and their colour shows the nature of samples (unspoiled, fermented, spoiled or both).

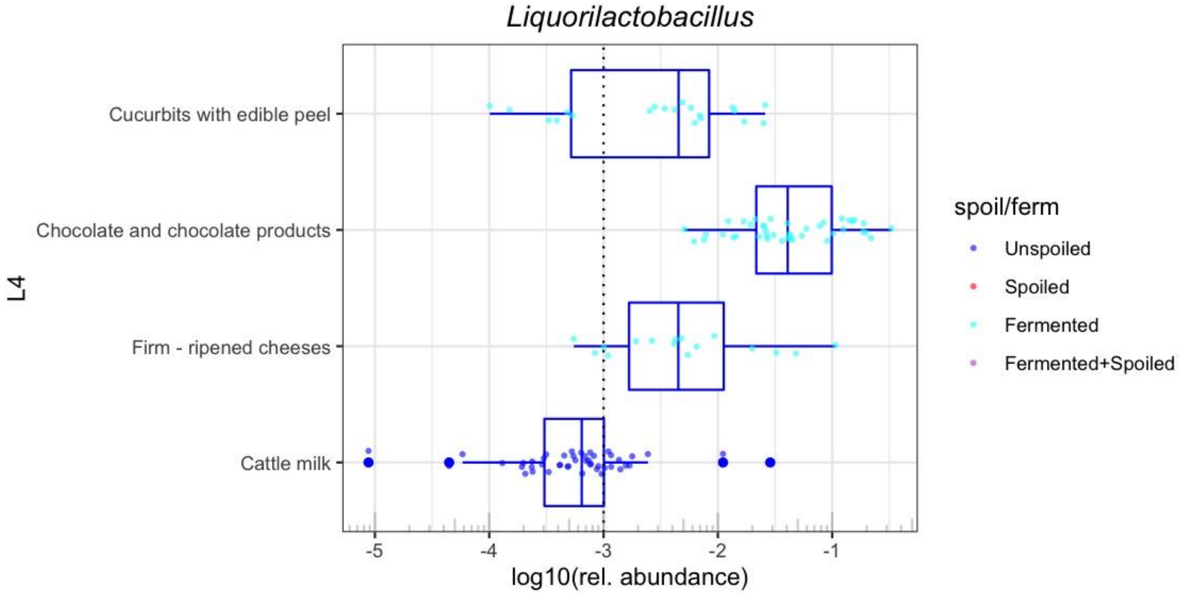

**Supplementary Figure 9**. Box and jitter showing the distribution of relative abundance of *Latilactobacillus* sequences (only sequences for the V1-V3 and V3-V4 regions with length >350 bp were selected) in food samples from FoodMicrobionet. The L4 level of the EFSA FoodEx2 classification is used for foods. The individual points show the abundance in samples and their colour shows the nature of samples (unspoiled, fermented, spoiled or both).

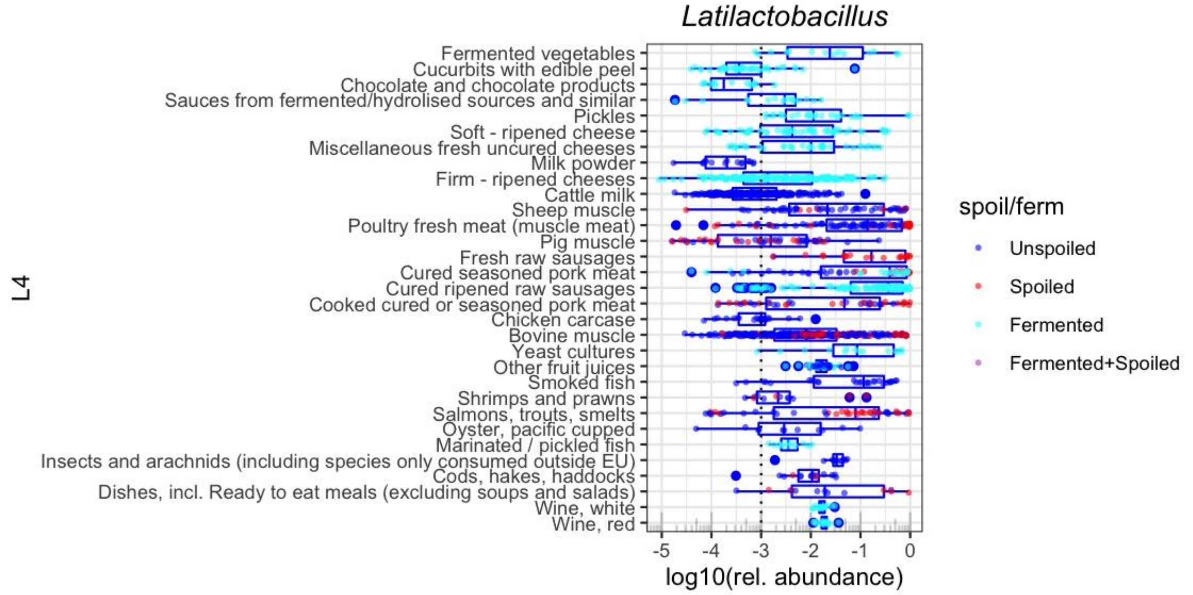


**Supplementary Figure 10**. Box and jitter showing the distribution of relative abundance of *Dellaglioa* sequences (only sequences for the V1-V3 and V3-V4 regions with length >350 bp were selected) in food samples from FoodMicrobionet. The L4 level of the EFSA FoodEx2 classification is used for foods. The individual points show the abundance in samples and their colour shows the nature of samples (unspoiled, fermented, spoiled or both).

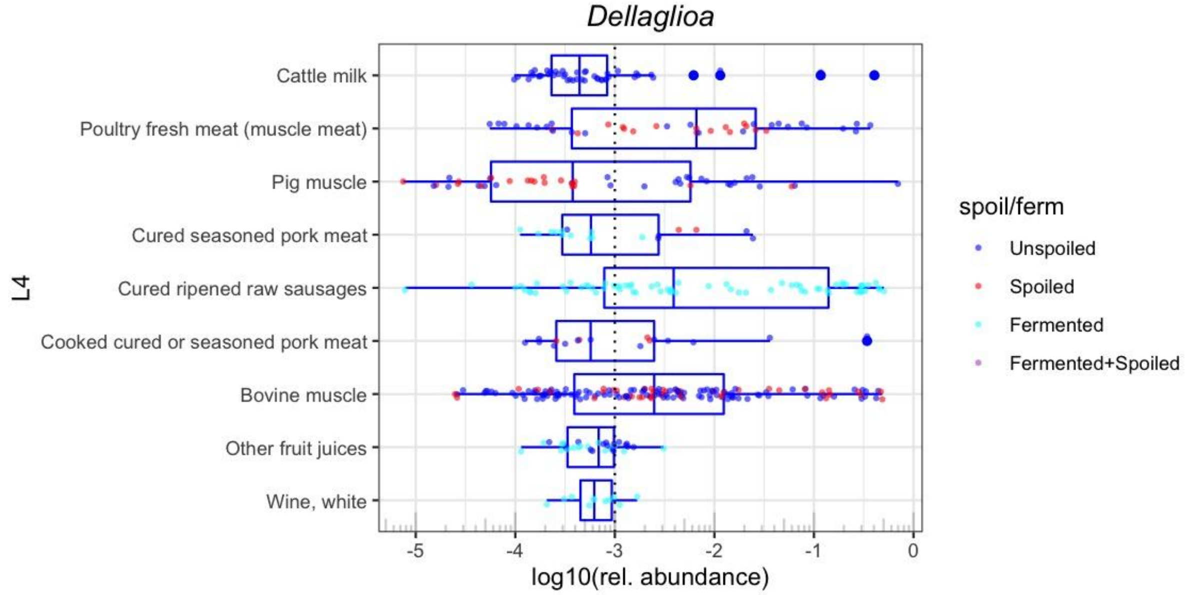

**Supplementary Figure 11**. Box and jitter showing the distribution of relative abundance of *Dellaglioa* sequences (only sequences for the V1-V3 and V3-V4 regions with length >350 bp were selected) in food samples from FoodMicrobionet. The L4 level of the EFSA FoodEx2 classification is used for foods. The individual points show the abundance in samples and their colour shows the nature of samples (unspoiled, fermented, spoiled or both).

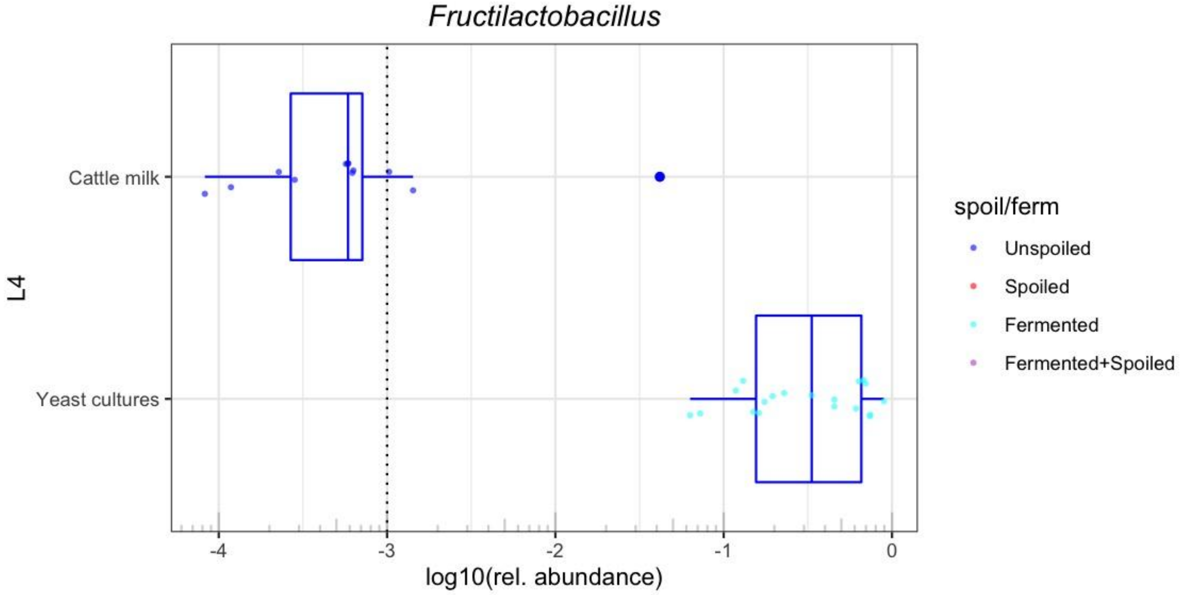


**Supplementary Figure 12**. Box and jitter showing the distribution of relative abundance of *Levilactobacillus* sequences (only sequences for the V1-V3 and V3-V4 regions with length >350 bp were selected) in food samples from FoodMicrobionet. The L4 level of the EFSA FoodEx2 classification is used for foods. The individual points show the abundance in samples and their colour shows the nature of samples (unspoiled, fermented, spoiled or both).

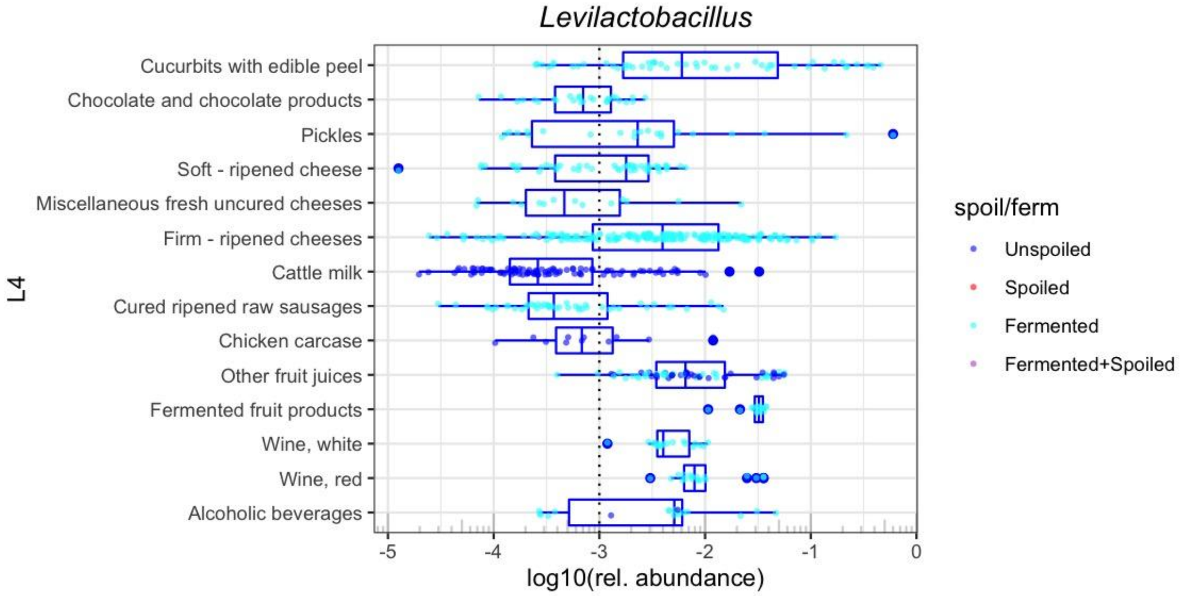

**Supplementary Figure 13**. Box and jitter showing the distribution of relative abundance of *Weissella* sequences (only sequences for the V1-V3 and V3-V4 regions with length >350 bp were selected) in food samples from FoodMicrobionet. The L4 level of the EFSA FoodEx2 classification is used for foods. The individual points show the abundance in samples and their colour shows the nature of samples (unspoiled, fermented, spoiled or both).

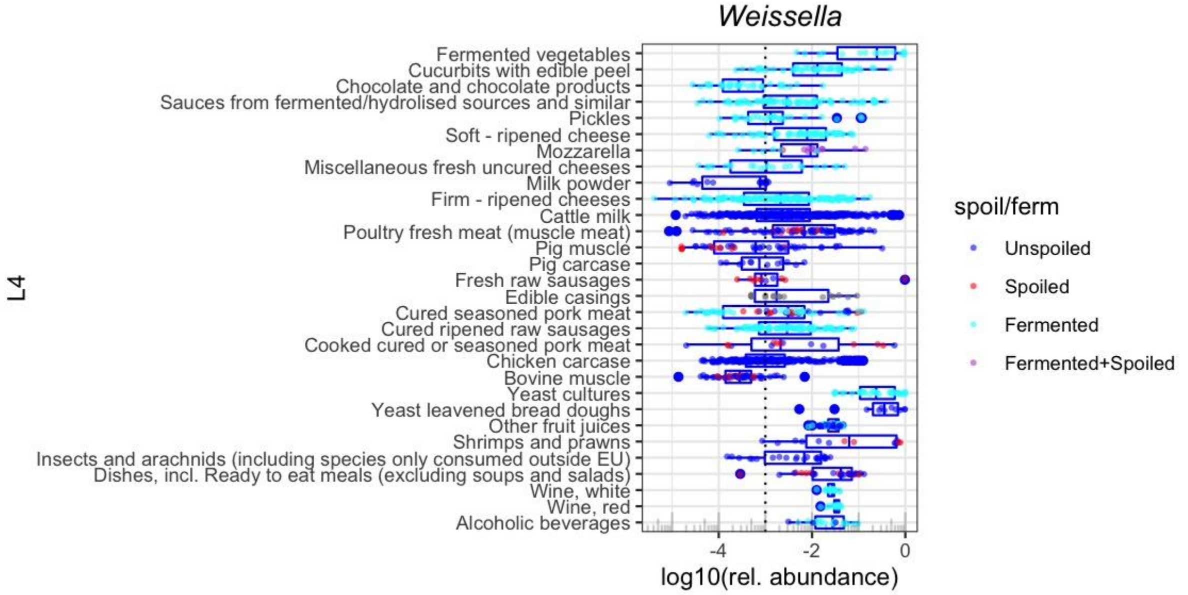


**Supplementary Figure 14**. Box and jitter showing the distribution of relative abundance of *Fructobacillus* sequences (only sequences for the V1-V3 and V3-V4 regions with length >350 bp were selected) in food samples from FoodMicrobionet. The L4 level of the EFSA FoodEx2 classification is used for foods. The individual points show the abundance in samples and their colour shows the nature of samples (unspoiled, fermented, spoiled or both).

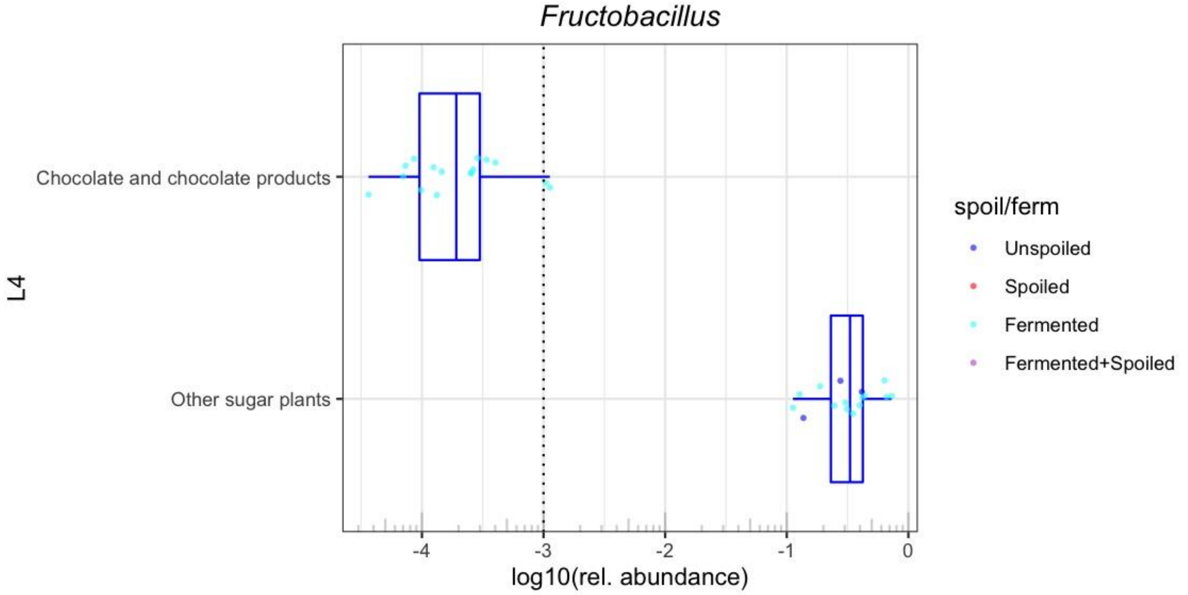

**Supplementary Figure 15**. Maximum likelihood phylogenetic tree showing the relationships between Amplicon Sequence Variants identified as belonging to the genus *Dellaglioa* (V1-V3 region). Reference sequences extracted from the SILVA v138.1 database, type strain sequences and outgroups are included. Further annotation includes color (food or food environment group where the sequences were found) and size (proportional to the maximum relative abundance of a given ASV).

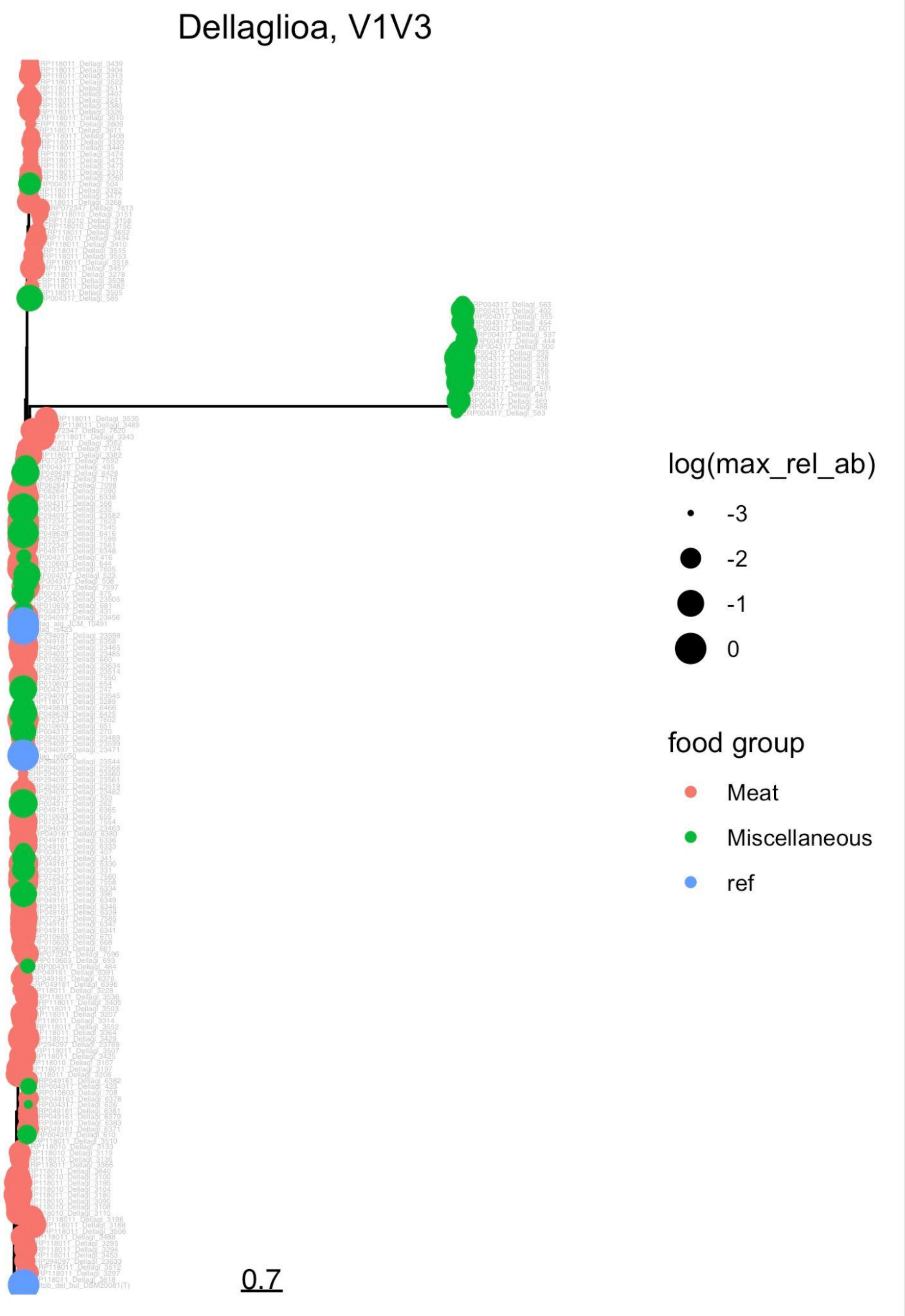

**Supplementary Figure 16**. Maximum likelihood phylogenetic tree showing the relationships between Amplicon Sequence Variants identified as belonging to the genus *Dellaglioa* (V3-V4 region). Reference sequences extracted from the SILVA v138.1 database, type strain sequences and outgroups are included. Further annotation includes color (food or food environment group where the sequences were found) and size (proportional to the maximum relative abundance of a given ASV).

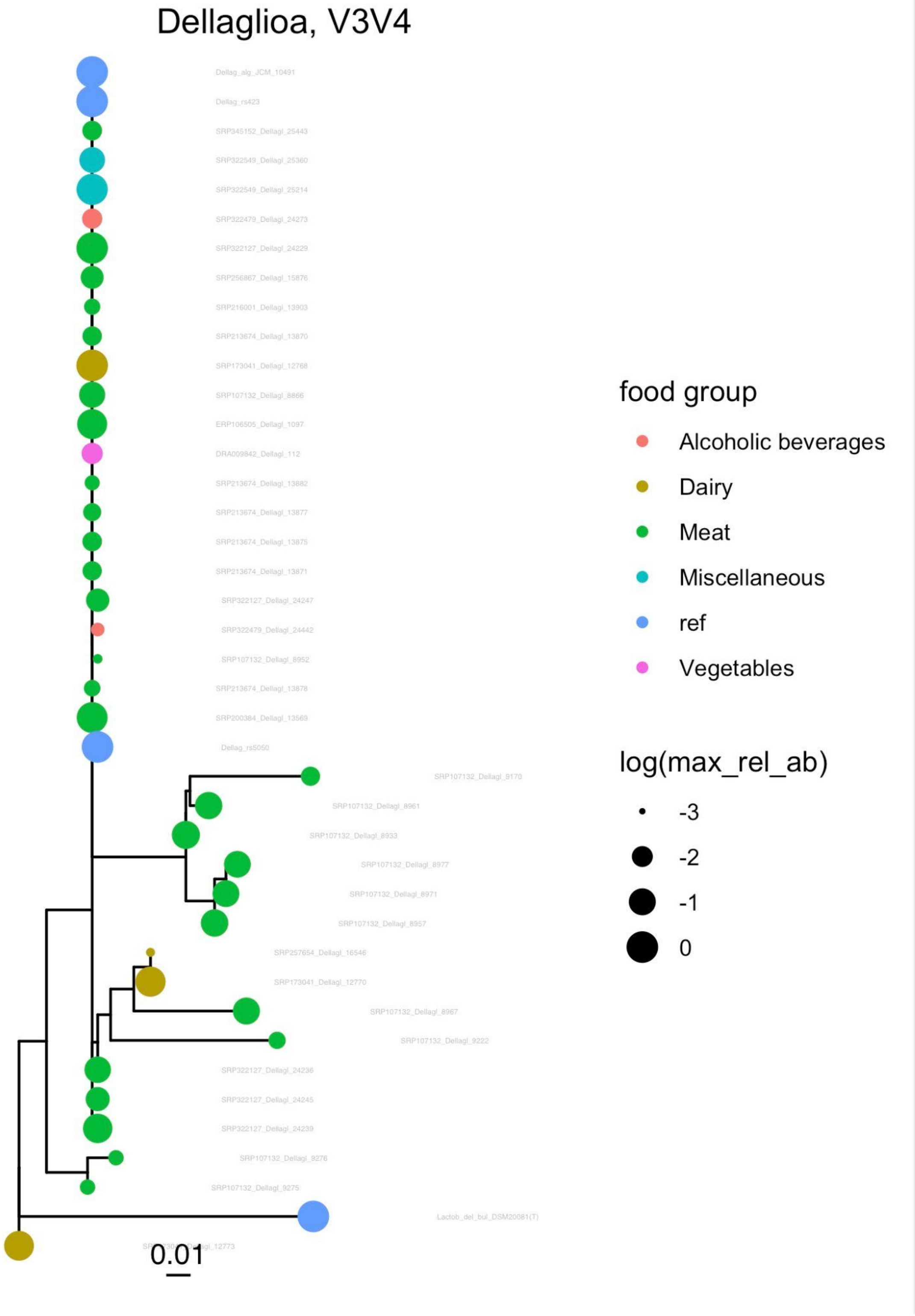

**Supplementary Figure 17**. Maximum likelihood phylogenetic tree showing the relationships between Amplicon Sequence Variants identified as belonging to the genus *Holzapfelia* (V1-V3 region). Reference sequences extracted from the SILVA v138.1 database, type strain sequences and outgroups are included. Further annotation includes color (food or food environment group where the sequences were found) and size (proportional to the maximum relative abundance of a given ASV).

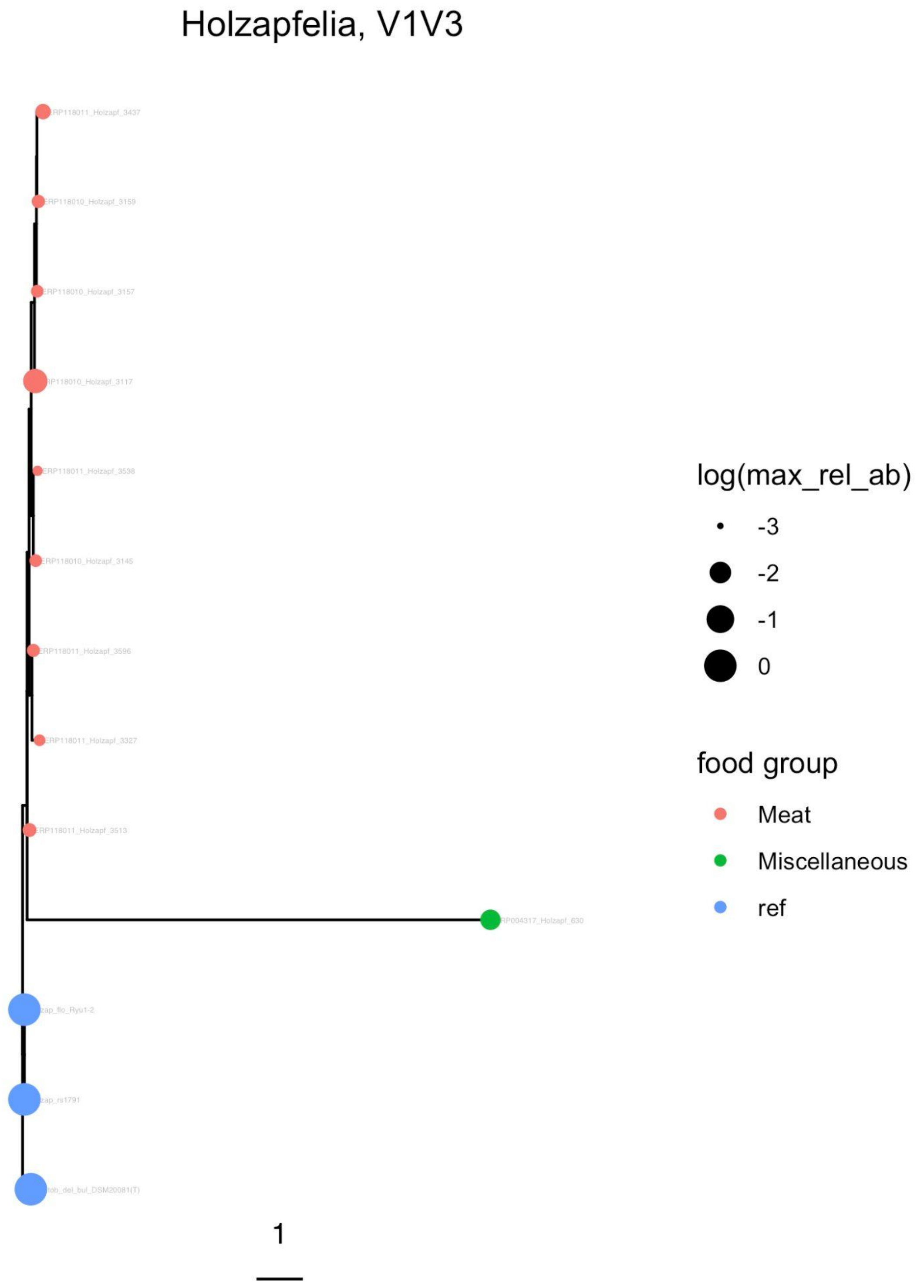

**Supplementary Figure 18**. Maximum likelihood phylogenetic tree showing the relationships between Amplicon Sequence Variants identified as belonging to the genus *Holzapfelia* (V3-V4 region). Reference sequences extracted from the SILVA v138.1 database, type strain sequences and outgroups are included. Further annotation includes color (food or food environment group where the sequences were found) and size (proportional to the maximum relative abundance of a given ASV).

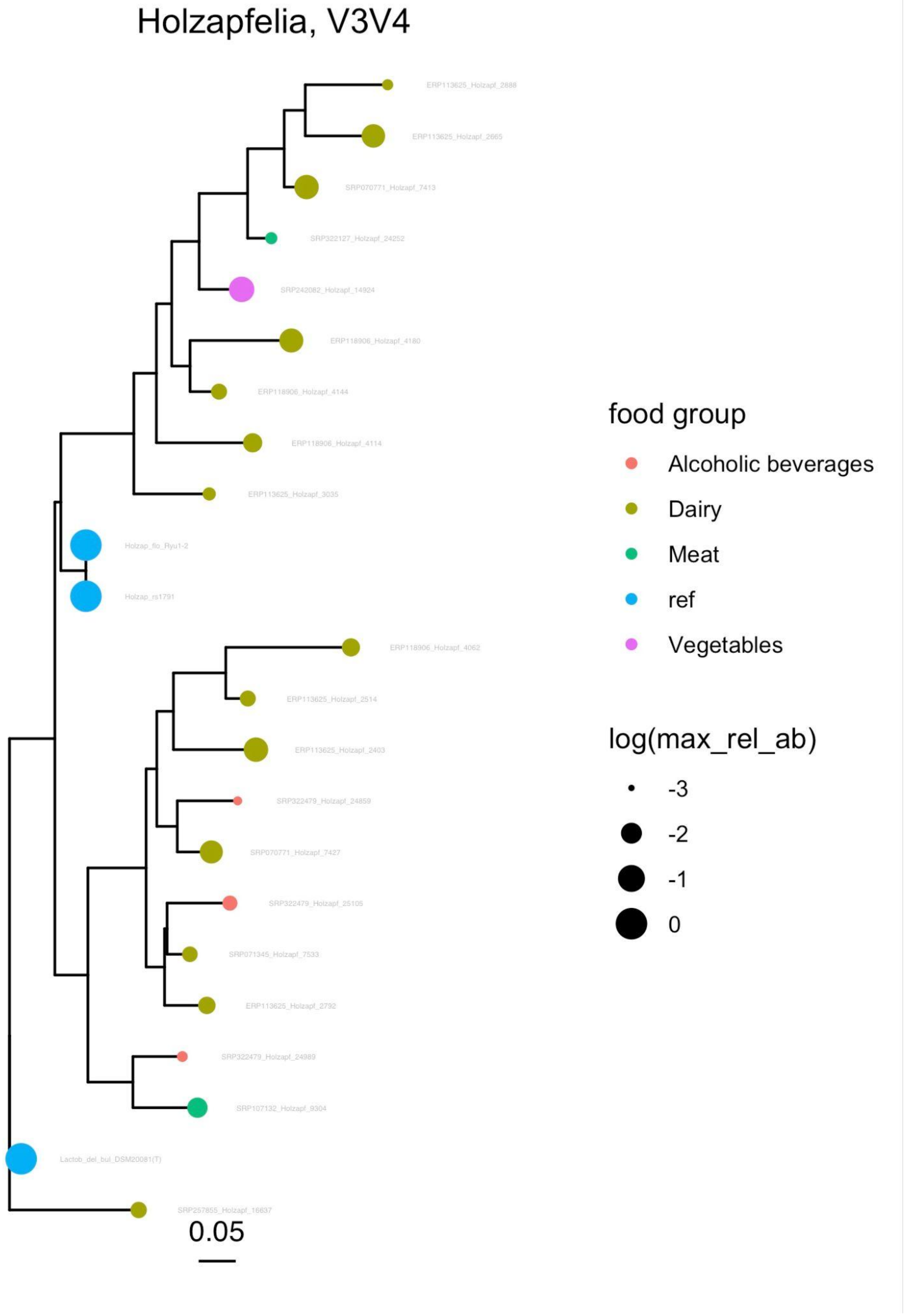

**Supplementary Figure 19**. Maximum likelihood phylogenetic tree showing the relationships between Amplicon Sequence Variants identified as belonging to the genus *Schleiferilactobacillus* (V1-V3 region). Reference sequences extracted from the SILVA v138.1 database, type strain sequences and outgroups are included. Further annotation includes color (food or food environment group where the sequences were found) and size (proportional to the maximum relative abundance of a given ASV).

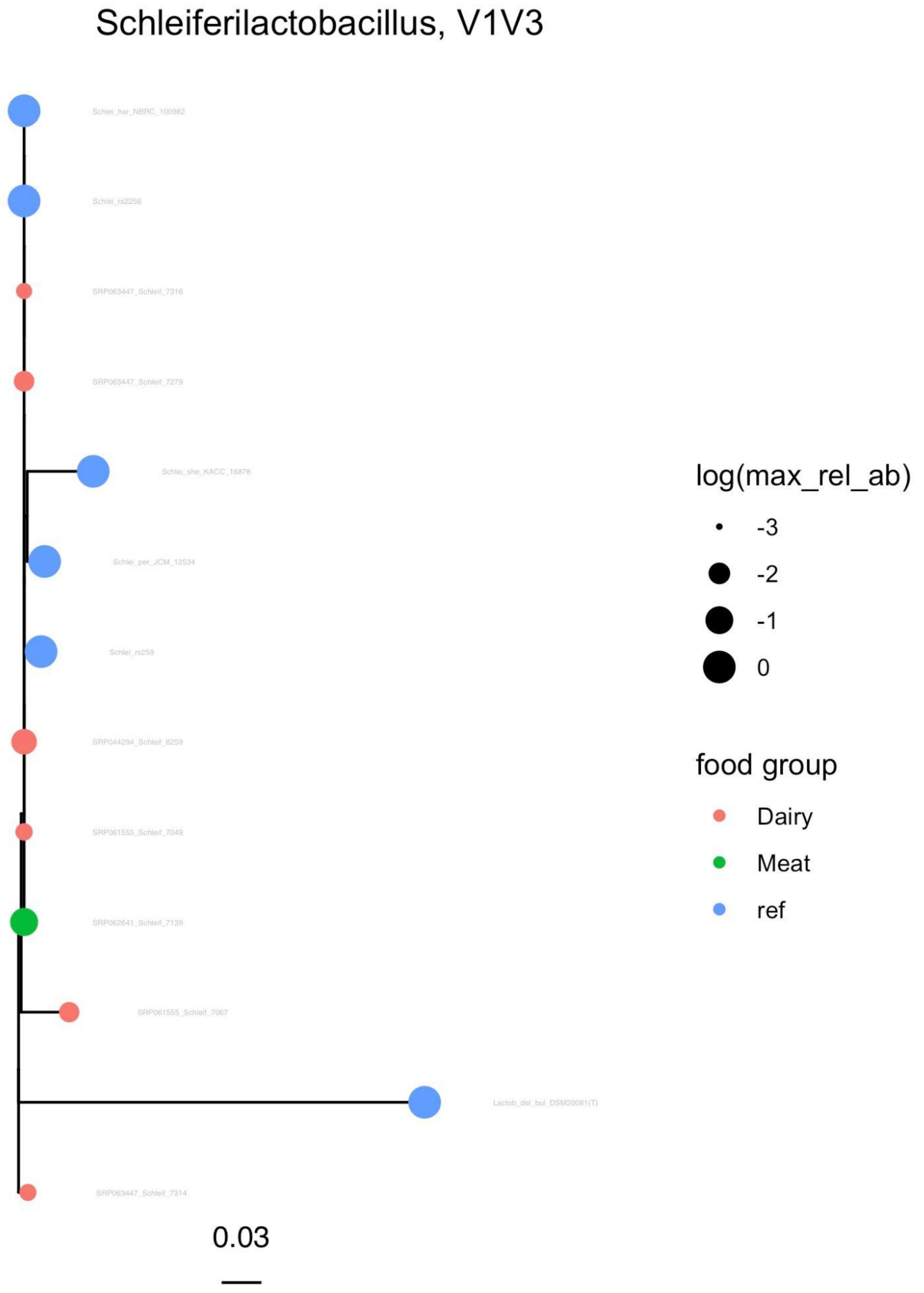

**Supplementary Figure 20**. Maximum likelihood phylogenetic tree showing the relationships between Amplicon Sequence Variants identified as belonging to the genus *Schleiferilactobacillus* (V3-V4 region). Reference sequences extracted from the SILVA v138.1 database, type strain sequences and outgroups are included. Further annotation includes color (food or food environment group where the sequences were found) and size (proportional to the maximum relative abundance of a given ASV).

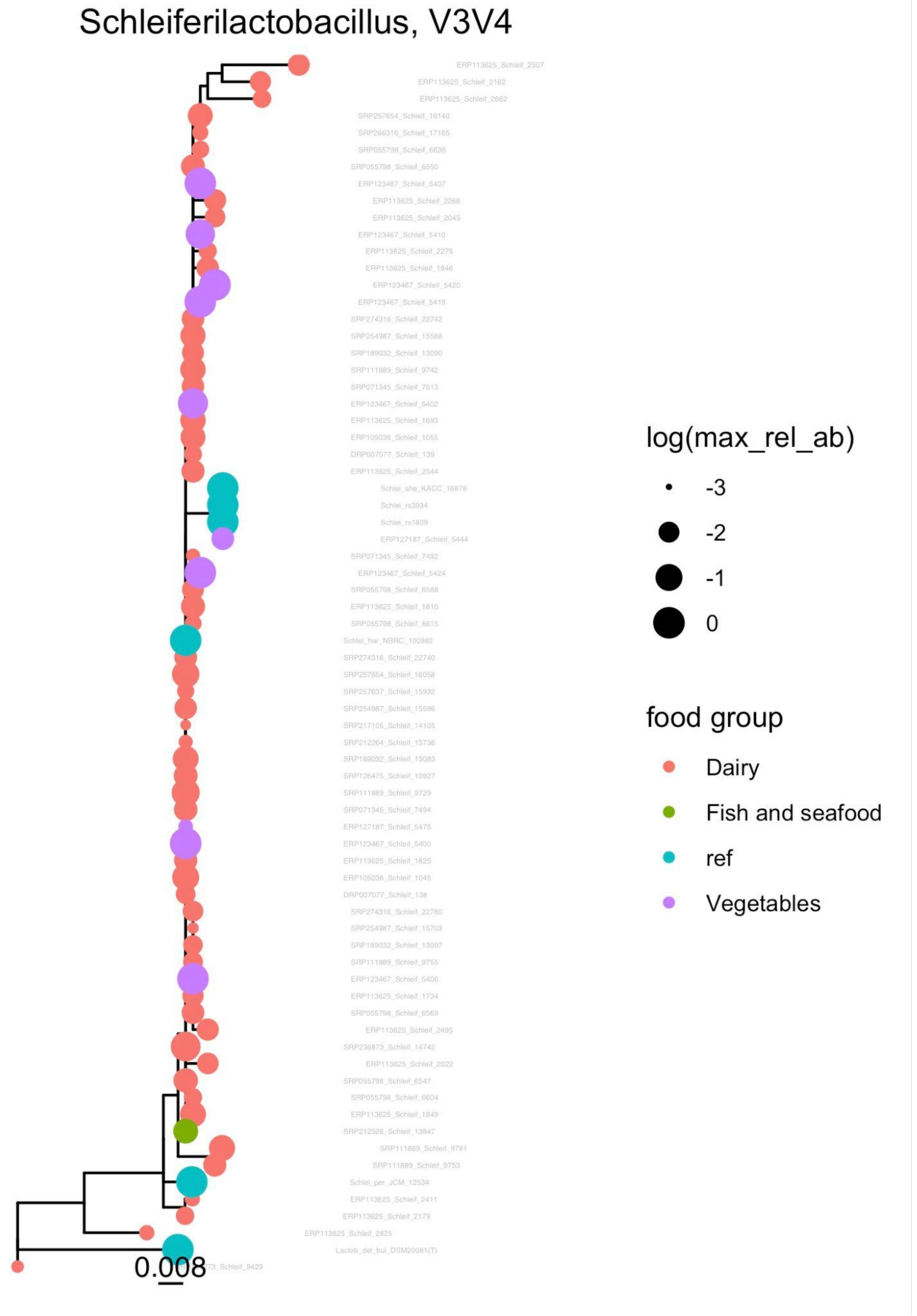
